# A helper NLR monitors catalytic perturbation of a vesicle trafficking regulator targeted by pathogen effectors

**DOI:** 10.64898/2026.10.06.757045

**Authors:** Enoch Lok Him Yuen, Nick Eilmann, Yijun Wang, Tolga O. Bozkurt

**Author notes:** Correspondence (T.O.B.).

## Abstract

Plants rely on vesicle trafficking for immunity, yet pathogens deploy effectors to reprogram these pathways. NLR immune receptors can monitor trafficking components, but how plants detect effector perturbation of the enzymatic regulators controlling these pathways remains unclear. Here, we show that TOPGAP, a Rab GTPase-activating protein and susceptibility factor, couples effector activity to immune signaling involving the helper NLR NRC4. PiE355 and TIKI, related effectors from two *Phytophthora* species, bind the N-terminal regulatory domain of TOPGAP and trigger cell death requiring TOPGAP and its C-terminal GAP activity. The TOPGAP catalytic domain alone is sufficient to activate NRC4-mediated cell death, which is abolished by catalytic inactivation and suppressed by the N-terminal regulatory domain that the effectors bind. These findings implicate deregulated TOPGAP activity in immune activation. We propose that NRC4-dependent surveillance enables plants to detect deregulation of a host enzyme that pathogens can co-opt to promote infection.

## Introduction

Plant cells rely on membrane trafficking to mount effective immune responses. Upon pathogen attack, immune-related secretory pathways are redirected to focus defense components at the site of pathogen invasion^1,2^. This focal secretion is particularly prominent during infection by filamentous pathogens that form haustoria. These specialized infection structures are enveloped by a host-derived extrahaustorial membrane (EHM), a key interface between plant and pathogen^3,4^.

As in other eukaryotes, secretory trafficking in plants is controlled by Rab GTPases, which switch between a GTP-bound active state and a GDP-bound inactive state. Guanine nucleotide exchange factors (GEFs) promote GDP–GTP exchange and Rab activation, whereas GTPase-activating proteins (GAPs) accelerate GTP hydrolysis and Rab inactivation^5–7^.

These secretory routes are a prime target for pathogen effectors. Interactome screens place vesicle trafficking among the most heavily targeted host processes, and effectors from distantly related pathogens converge on the same components, including the Rab GTPases themselves^1,8,9^. We recently identified the host Rab GTPase-activating protein TOPGAP as a susceptibility factor for *Phytophthora infestans* targeted by a conserved family of oomycete effectors^10^. TOPGAP comprises an N-terminal Rab-binding domain (RBD) and a C-terminal Tre2/Bub2/Cdc16 (TBC) domain that harbors its GAP catalytic activity. TIKI from *Phytophthora palmivora* and its *P. infestans* homologs PiE354 and PiE355 associate with TOPGAP through its RBD. PiE354 exploits TOPGAP’s catalytic activity on Rab8a, depleting this GTPase from the plasma membrane and diverting defense-related secretion away from the pathogen interface^10^. TOPGAP thus provides a host enzymatic activity that pathogens can co-opt to suppress immunity and promote infection.

Plants counter effectors that target critical defense-related pathways by deploying intracellular nucleotide-binding leucine-rich repeat immune receptors (NLRs)^11–13^. Some NLRs activate immunity by binding effectors directly^14,15^, whereas others monitor the host proteins that effectors act upon and are activated by their modification, cleavage or degradation^16–18^. In solanaceous plants, many sensor NLRs cannot signal alone and depend on a partially redundant network of helper NLRs, the NRCs (NLR required for cell death), to execute immunity^19,20^. Activated NRCs oligomerize into multimeric resistosomes and execute cell death through a shared N-terminal MADA motif^21–25^. Among them, NRC4 accumulates at the EHM during *Phytophthora* infection, localizing this helper NLR to a critical site of defense-related membrane trafficking^26^.

The convergence of pathogen effectors on defense-related membrane trafficking raises the question of how plants safeguard its enzymatic regulators. Although NLR-mediated surveillance has been linked to individual trafficking components^27,28^, how plants detect perturbation of the enzymes controlling these pathways remains poorly understood. TIKI, PiE355 and PiE354 all associate with TOPGAP, yet only TIKI and PiE355 elicit cell death in *Nicotiana benthamiana*^10^. Their shared target but contrasting outcomes raise the possibility that immune activation depends on how these effectors perturb TOPGAP function. Whether the GAP activity exploited to promote susceptibility also contributes to effector-triggered immune activation remains unresolved.

Here we show that TOPGAP, a susceptibility factor exploited to suppress defense-related trafficking, also contributes to effector-triggered immune activation involving the helper NLR NRC4. TOPGAP silencing reduces cell death elicited by PiE355 and TIKI, and TOPGAP-mediated enhancement of these responses requires both GAP activity and effector association. Genetic complementation identifies NRC4 as the principal NRC helper supporting both responses. The isolated TBC domain induces cell death in the absence of effectors; this response requires GAP activity, is suppressed by the N-terminal RBD and is attenuated by loss of NRC4. Together, these findings support a model in which deregulated TOPGAP activity provides a cue for immune surveillance that signals through NRC4.

## Results

### 1. PiE355-triggered cell death depends on TOPGAP activity and effector engagement

The *Phytophthora infestans* RXLR effectors PiE354 and PiE355 both target TOPGAP, a host Rab GTPase-activating protein and susceptibility factor. We previously showed that PiE354 exploits TOPGAP to redirect defense-related trafficking and promote infection^10^. Although both effectors associate with TOPGAP, PiE355 elicits cell death in *Nicotiana benthamiana*, whereas PiE354 does not. We therefore investigated whether PiE355-triggered cell death depends on TOPGAP and its catalytic function.

RNAi-mediated silencing of TOPGAP significantly reduced PiE355-triggered cell death compared with the control construct targeting β-glucuronidase (RNAi:GUS; Figs. 1A, B). Neither silencing construct induced cell death when co-expressed with an empty vector (EV). Conversely, co-expression of TOPGAP with PiE355 significantly enhanced cell death relative to the EV control (Figs. 1C, D). This enhancement depended on TOPGAP catalytic competence: the GAP-inactive mutant TOPGAP^GAP^ failed to enhance the response and instead significantly reduced it relative to the EV control. By contrast, PiE354 did not elicit visible cell death when co-expressed with TOPGAP, TOPGAP^GAP^ or EV (Fig. S1). These results identify TOPGAP as a contributor to PiE355-triggered cell death and implicate its GAP activity in this response.

**Figure 1.**
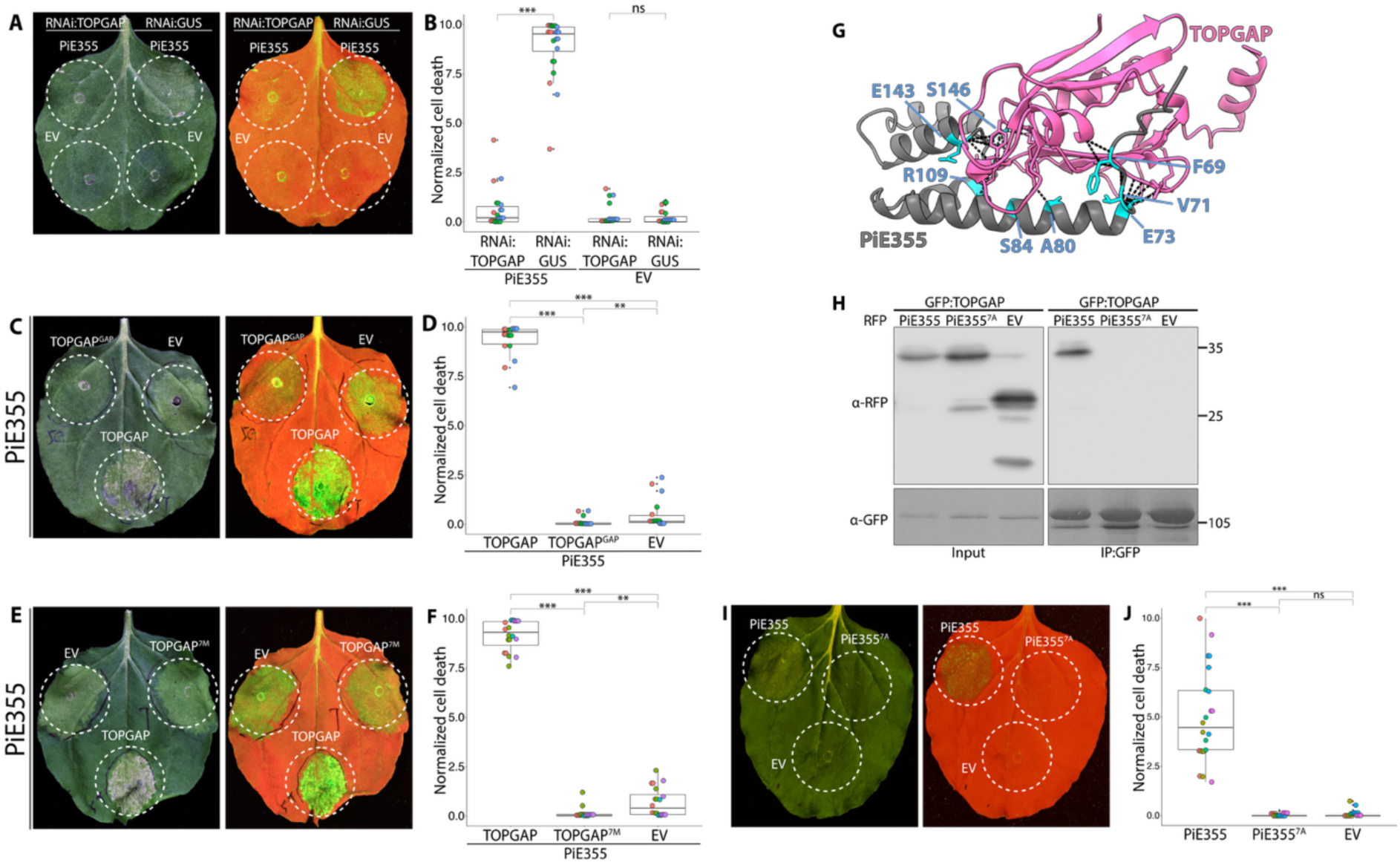
TOPGAP catalytic activity and effector association contribute to PiE355-triggered cell death. (A–B) Silencing of TOPGAP significantly reduces cell death induced by PiE355. (A) Representative images of *N. benthamiana* leaves infiltrated with PiE355 or EV control, along with RNAi:TOPGAP or the silencing control RNAi:GUS. Daylight and UV images were captured at 4 dpi, and cell death was quantified. (B) Box-and-dot plot showing that silencing TOPGAP significantly reduces cell death triggered by PiE355 (0.59, n = 24), compared to the GUS-silencing control (8.95, n = 24). EV controls do not induce visible cell death. (C–D) Overexpression of TOPGAP increases cell death induced by PiE355, dependent on its GAP activity. (C) Representative images of *N. benthamiana* leaves infiltrated with PiE355, with TOPGAP, TOPGAP^GAP^, or EV. Daylight and UV images were captured at 4 dpi, and cell death was quantified. (D) Box-and-dot plot showing that overexpression of TOPGAP significantly increases cell death triggered by PiE355 (9.36, n = 18), compared to the EV control (0.52, n = 18), while the GAP mutant significantly reduces cell death triggered by PiE355 (0.12, n = 18). (E–F) Overexpression of TOPGAP increases cell death induced by PiE355 dependent on their interaction. (E) Representative images of *N. benthamiana* leaves infiltrated with PiE355, with TOPGAP, TOPGAP^7M^, or EV. Daylight and UV images were captured at 4 dpi, and cell death was quantified. (F) Box-and-dot plot showing that overexpression of TOPGAP significantly increases cell death triggered by PiE355 (9.18, n = 20), compared to the EV control (0.70, n = 20), while the mutant TOPGAP^7M^ significantly reduces cell death triggered by PiE355 (0.13, n = 20). (G) AF2-M–predicted model of PiE355 (gray) targeting the RBD of TOPGAP (pink), depicting the interacting residues on PiE355. The bonds between PiE355 and TOPGAP are predicted by PyMOL with a distance of 3 Å. PiE355 interacts with TOPGAP using eight key residues on PiE355, colored as cyan. (H) Substitution of seven predicted interface residues abolishes detectable association of PiE355 with TOPGAP. GFP:TOPGAP was transiently co-expressed with either RFP:PiE355, RFP:PiE355^7A^, or RFP:EV. (I–J) The interaction-deficient mutant PiE355^7A^ does not cause visible cell death. (I) Representative images of *N. benthamiana* leaves infiltrated with PiE355, PiE355^7A^ or EV control. Daylight and UV images were captured at 4 dpi, and cell death was quantified. (J) Box-and-dot plot showing that while PiE355 causes cell death (5.00, n = 22), PiE355^7A^ and the EV control do not cause visible cell death. Each color represents an independent biological replicate. Statistical comparisons were performed using Mann–Whitney U tests with Bonferroni correction in R. **P < 0.01; ***P < 0.001. ns, not significant.

To assess whether TOPGAP broadly modulates effector-triggered cell death, we examined responses to two additional RXLR effectors. AVRblb2 is recognized by the sensor NLR Rpi-blb2, triggering hypersensitive cell death^29–31^. Neither wild-type TOPGAP nor the GAP-inactive mutant TOPGAP^GAP^ significantly altered cell death triggered by co-expression of AVRblb2 and Rpi-blb2 (Fig. S2A). Similarly, neither construct significantly affected cell death elicited by RxLR24, a *Phytophthora* effector that interacts with Rab GTPases involved in vesicle trafficking^32^ (Fig. S2B). These results indicate that the enhancement of PiE355-triggered cell death by TOPGAP is not a general effect on the other effector-triggered responses tested.

We next examined whether effector–target interaction contributes to the response. Unlike wild-type TOPGAP, the interaction-deficient mutant TOPGAP^7M^ failed to enhance PiE355-triggered cell death^10^ (Figs. 1E, F), indicating that TOPGAP-mediated enhancement requires an interaction-competent host target.

To further test whether association with TOPGAP is required for PiE355-triggered cell death, we used AlphaFold-Multimer to guide mutagenesis of the predicted interaction interface. The model identified eight candidate interface residues with high model confidence (predicted aligned error, PAE < 5 Å) in PiE355, seven of which were substituted with alanine to generate PiE355^7A^ (Fig. 1G; Fig. S3). Wild-type PiE355 co-immunoprecipitated with TOPGAP, whereas PiE355^7A^ showed no detectable association (Fig. 1H). Loss of detectable association was accompanied by loss of cell-death activity: PiE355^7A^ failed to elicit detectable cell death under conditions in which wild-type PiE355 was active (Figs. 1I, J).

Together, these findings identify TOPGAP as a mediator of PiE355-triggered cell death, with both its catalytic activity and interaction with the effector contributing to this response.

### 2. The helper NLR NRC4 mediates PiE355-triggered hypersensitive cell death

These findings raised the possibility that PiE355 targeting of TOPGAP activates immune surveillance of the host protein. We therefore asked whether PiE355-triggered cell death engages NLR-mediated immune signaling. Re-analysis of the TOPGAP immunoprecipitation-mass spectrometry (IP-MS) dataset from our previous study identified the helper NLR NRC4 as a candidate interactor, pointing to a possible role in PiE355-triggered cell death (Table S1)^10^.

We first examined whether NRC4 could potentiate PiE355-triggered cell death in wild-type *Nicotiana benthamiana*. Co-expression of NRC4 significantly enhanced the response to PiE355 compared with the EV control, whereas NRC4 did not induce detectable cell death when expressed with PiE354 or EV (Figs. 2A, B). Co-expression of the sensor NLR Rpi-blb2 and its cognate effector AVRblb2 elicited strong hypersensitive response (HR) cell death and served as a positive control for an established NRC4-dependent immune response^20^.

**Figure 2.**
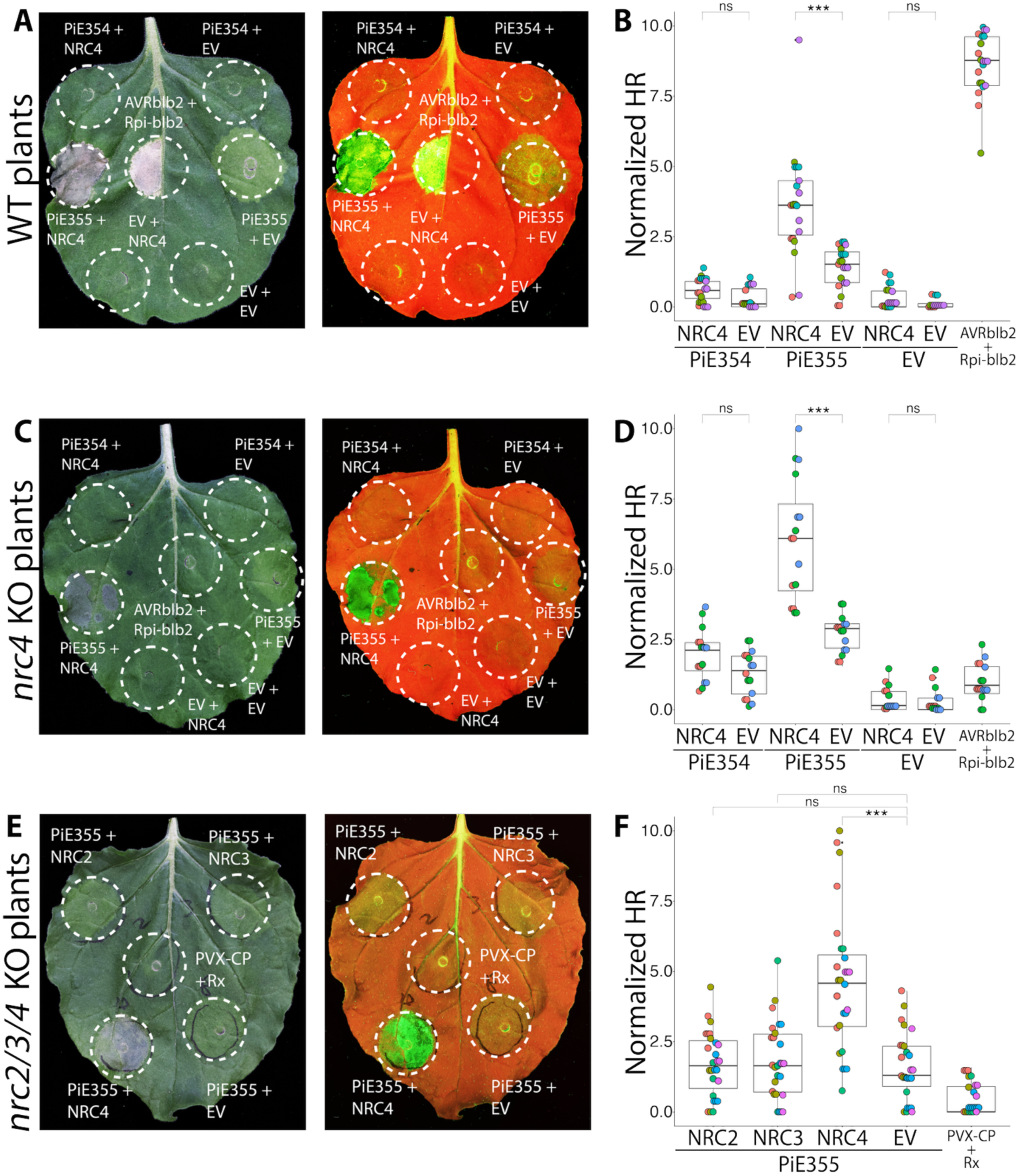
NRC4 mediates robust PiE355-triggered hypersensitive cell death. (A–B) Overexpression of NRC4 enhances PiE355-triggered HR in WT plants. (A) Representative images of WT *N. benthamiana* leaves infiltrated with PiE354, PiE355 or EV control, together with NRC4 or EV. Daylight and UV images were captured at 4 dpi and HR was quantified. (B) Box-and-dot plot showing that NRC4 overexpression significantly increases HR triggered by PiE355 (3.63, n = 21) compared with EV control (1.42, n = 21). (C–D) Genetic loss and complementation establish a contribution of NRC4 to PiE355-triggered HR. (C) Representative images of *nrc4* KO plants infiltrated with PiE354, PiE355 or EV, together with NRC4 or EV. In *nrc4* KO plants, PiE355-induced HR is reduced compared with WT, and reintroduction of NRC4 restores robust HR. Images were captured at 4 dpi and HR was quantified. (D) Box-and-dot plot showing that PiE355-induced HR is attenuated in *nrc4* KO plants expressing EV (2.71, n = 16), whereas complementation with NRC4 restores and significantly enhances HR (6.04, n = 16). NRC4 expression does not significantly alter the response to PiE354 or to the EV control. Co-expression of AVRblb2 and Rpi-blb2 was used as an HR control. (E–F) NRC2 and NRC3 do not substitute for NRC4 in supporting PiE355-triggered HR. (E) Representative images of *nrc2/3/4* KO plants infiltrated with PiE355 together with NRC2, NRC3, NRC4 or EV. Images were captured at 4 dpi, and HR was quantified. (F) Box-and-dot plot showing that only NRC4 expression significantly enhances PiE355-triggered HR (4.62, n = 27) compared with EV (1.6, n = 27), whereas NRC2 (1.72, n = 27) and NRC3 (1.81, n = 27) do not significantly alter the response. Co-expression of PVX-CP and Rx was used as an HR control. Each color represents an independent biological replicate. Statistical comparisons were performed using Mann–Whitney U tests with Bonferroni correction in R. ***P < 0.001. ns, not significant.

To investigate the contribution of NRC4 genetically, we expressed PiE355 in plants lacking NRC4a/b (*nrc4* KO plants). PiE355 triggered only weak cell death in *nrc4* KO plants compared with wild-type plants, though the response was not fully abolished. Reintroduction of NRC4 significantly enhanced the response, producing robust tissue collapse (Figs. 2C, D). Neither PiE354 nor EV elicited detectable cell death upon NRC4 expression. The AVRblb2–Rpi-blb2 pair failed to elicit visible HR in uncomplemented *nrc4* KO plants, consistent with disruption of NRC4-dependent signaling.

We next asked whether other NRC helpers could substitute for NRC4. In plants lacking NRC2, NRC3 and NRC4 (*nrc2/3/4* KO plants), co-expression of NRC4 significantly enhanced PiE355-triggered cell death, whereas NRC2 and NRC3 did not significantly alter the response relative to EV (Figs. 2E, F). The sensor NLR Rx and its elicitor, potato virus X coat protein (PVX-CP), failed to trigger visible HR in uncomplemented *nrc2/3/4* KO plants, consistent with the established dependence of this response on NRC helpers^33^. These results indicate that NRC2 and NRC3 do not substitute for NRC4 in promoting PiE355-triggered cell death.

To test whether PiE355-triggered HR requires the execution activity of the NRC4 resistosome, we used NRC4^L9E^, a MADA-motif mutant that compromises NRC4-mediated calcium influx and cell death while retaining the capacity for resistosome assembly^22,34,35^. We co-expressed PiE355 with either wild-type NRC4 or NRC4^L9E^ in *nrc4* KO plants. Wild-type NRC4 supported robust cell death, whereas the response supported by NRC4^L9E^ was significantly weaker (Fig. S4). Neither NRC4 variant induced detectable cell death when co-expressed with EV. These results show that robust PiE355-triggered HR depends on the MADA-mediated execution function of NRC4, supporting the involvement of NRC4 resistosome signaling in this response.

Together, these findings identify NRC4 as the principal NRC helper supporting PiE355-triggered hypersensitive cell death and connect this response to its MADA-dependent execution activity. Combined with the dependence on TOPGAP described above, these results support a model in which PiE355 targeting of TOPGAP engages NRC4-dependent immune signaling.

### 3. The TOPGAP catalytic domain triggers GAP-dependent cell death promoted by NRC4

TOPGAP enhanced PiE355-triggered cell death in a manner requiring its GAP catalytic activity, implicating its enzymatic function in this response. The C-terminal TBC domain of TOPGAP contains the conserved GAP catalytic residues R446 and Q483 (Fig. 3A; Fig. S5). We previously showed that expression of the isolated C-terminal TBC domain fragment of TOPGAP (TBCDF) induces cell death in *Nicotiana benthamiana*, whereas full-length TOPGAP and its N-terminal Rab-binding domain fragment (RBDF) do not^10^. This contrast raised the possibility that the N-terminal region restrains the cell-death activity of the catalytic domain. We therefore asked whether GAP activity and NRC4 contribute to TBCDF-triggered cell death, and whether the N-terminal Rab-binding domain suppresses this response.

**Figure 3.**
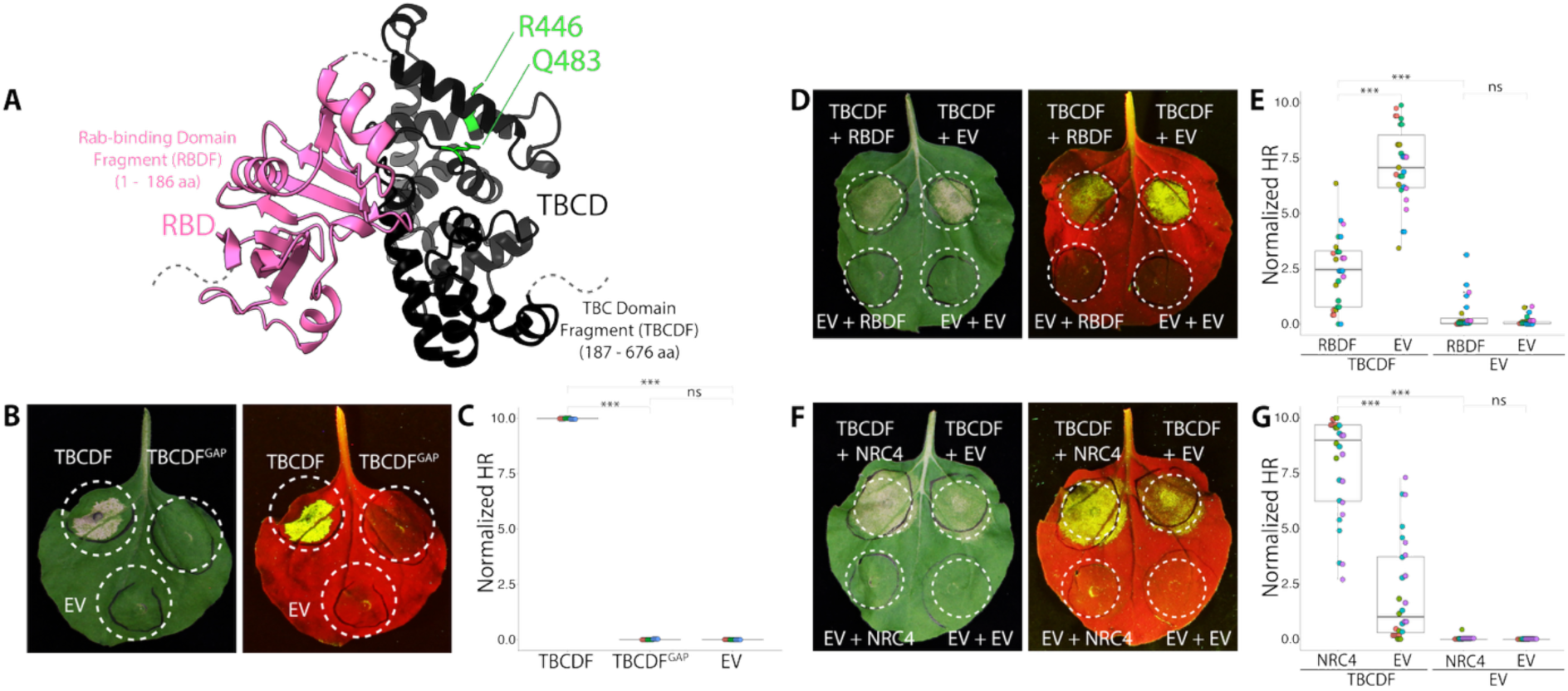
The TOPGAP catalytic domain triggers GAP-dependent cell death promoted by NRC4. (A) AlphaFold 3 (AF3)-predicted structure of full-length TOPGAP. The N-terminal Rab-binding domain (RBD; residues 1–186) is shown in pink and the C-terminal TBC domain (TBCD; residues 187–676) in black. The conserved catalytic residues R446 and Q483 are highlighted in green. (B–C) Overexpression of TBCDF causes HR in a GAP activity-dependent manner. (B) Representative images of *N. benthamiana* leaves infiltrated with TBCDF, TBCDF^GAP^, or EV. Daylight and UV images were captured at 4 dpi, and HR was quantified. (C) Box-and-dot plot showing that overexpression of TBCDF causes HR (9.99, n = 12), whereas the GAP-activity mutant TBCDF^GAP^ and the EV control do not elicit visible cell death. (D–E) Overexpression of RBDF reduces HR caused by TBCDF expression. (D) Representative images of *N. benthamiana* leaves infiltrated with TBCDF, or EV control, with RBDF, or EV control. Daylight and UV images were captured at 4 dpi, and HR was quantified. (E) Box-and-dot plot showing that overexpression of RBDF reduces HR caused by TBCDF expression (2.34, n = 27), compared to the EV control (7.17, n = 27), whereas the EV controls do not cause visible cell death. (F–G) Overexpression of NRC4 increases HR caused by TBCDF expression. (F) Representative images of *N. benthamiana* leaves infiltrated with TBCDF, or EV control, with NRC4, or EV control. Daylight and UV images were captured at 4 dpi, and HR was quantified. (G) Box-and-dot plot showing that overexpression of NRC4 increases HR caused by TBCDF expression (7.91, n = 28), compared to the EV control (2.07, n = 28), while the EV controls for TBCDF do not cause visible cell death. Each color represents an independent biological replicate. Statistical comparisons were performed using Mann–Whitney U tests with Bonferroni correction in R. ***P < 0.001. ns, not significant.

To test whether TBCDF-triggered cell death requires catalytic activity, we substituted the conserved catalytic arginine and glutamine residues with alanine to generate the GAP-inactive mutant TBCDFᴳᴬᴾ. Wild-type TBCDF elicited strong cell death, whereas TBCDFᴳᴬᴾ and the EV control did not (Figs. 3B, C). Immunoblotting confirmed stable expression of the mutant protein *in planta* (Fig. S6). These results establish that the cell-death activity of the isolated TBC domain depends on its GAP catalytic function.

The absence of cell death upon expression of full-length TOPGAP suggested that the N-terminal region might restrain the response elicited by the catalytic domain. Since both domains associate with Rab8a^10^, we reasoned that the N-terminal Rab-binding domain could influence substrate access to the catalytic domain. To test whether the N-terminal region can suppress TBCDF-triggered cell death, we co-expressed RBDF with TBCDF in wild-type plants. RBDF significantly reduced the response compared with EV, whereas RBDF alone did not induce detectable cell death (Figs. 3D, E). TBCDF accumulated to comparable levels when co-expressed with RBDF or EV (Fig. S7). Thus, the N-terminal fragment can suppress GAP-dependent cell death when supplied *in trans*.

We next asked whether TBCDF-triggered cell death engages NRC4, the helper NLR implicated in the response to PiE355. Co-expression of NRC4 significantly enhanced TBCDF-triggered cell death compared with EV, whereas NRC4 did not induce detectable cell death in the absence of TBCDF (Figs. 3F, G). Immunoblotting showed comparable TBCDF accumulation with NRC4 or EV (Fig. S8), indicating that the enhanced response was not associated with increased TBCDF abundance.

To test the genetic contribution of NRC4, we expressed full-length TOPGAP, RBDF, TBCDF or EV in wild-type and *nrc4* KO plants. Full-length TOPGAP, RBDF and EV did not elicit visible cell death in either genotype. TBCDF induced cell death in both backgrounds, but the response was weaker in *nrc4* KO plants (Figs. S9A, B). Together with the enhancement by NRC4 co-expression, this attenuation indicates that NRC4 contributes to the cell death response elicited by the TOPGAP catalytic domain.

Together, these findings show that the isolated TOPGAP catalytic domain triggers GAP-dependent cell death that is restrained by the N-terminal region and promoted by NRC4. These results support a model in which deregulation of TOPGAP activity engages NRC4-mediated immune signaling, providing a potential basis for surveillance of this effector-targeted host protein.

### 4. The *P. palmivora* effector TIKI engages the TOPGAP–NRC4 immune signaling module

We next asked whether the TOPGAP–NRC4 module also contributes to cell death triggered by TIKI, a PiE355 homolog from *Phytophthora palmivora* that shares only 50% amino acid identity across the effector domain, yet associates with TOPGAP and induces strong cell death in *N. benthamiana*^10^. This enabled us to test the contribution of the module to responses elicited by sequence-divergent effectors from different *Phytophthora* species.

We first examined the contribution of TOPGAP to TIKI-triggered cell death. Silencing TOPGAP significantly reduced the response compared with the RNAi:GUS control, whereas neither silencing construct induced detectable cell death when co-expressed with EV (Figs. 4A, B). Conversely, co-expression of wild-type TOPGAP significantly enhanced TIKI-triggered cell death. The GAP-inactive mutant TOPGAPᴳᴬᴾ failed to enhance the response and instead significantly suppressed it relative to EV (Figs. 4C, D). These results identify TOPGAP as a mediator of TIKI-triggered cell death and establish that its catalytic activity is required for enhancement of this response.

**Figure 4.**
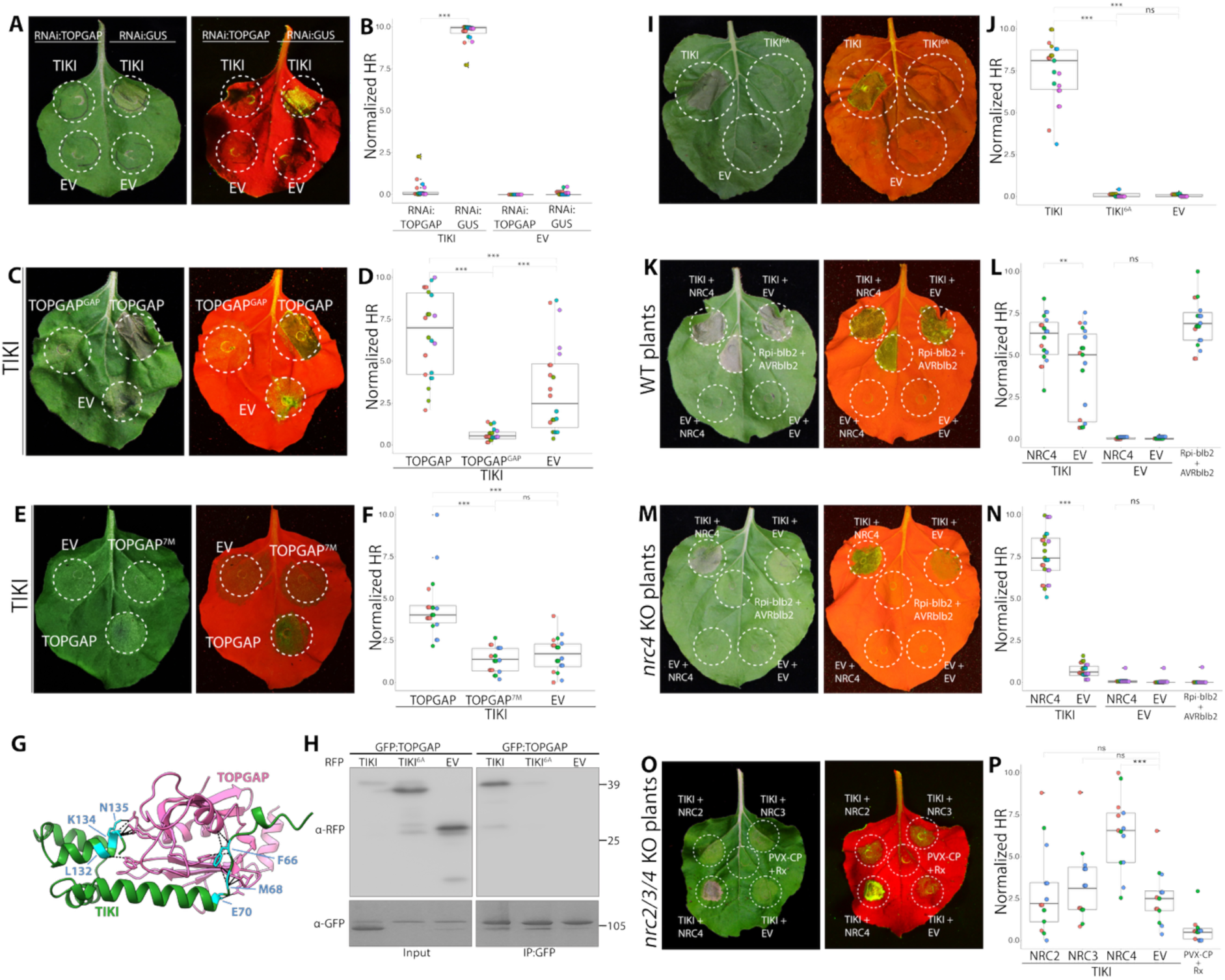
TOPGAP and NRC4 contribute to hypersensitive cell death triggered by *P. palmivora* effector TIKI. (A–B) Silencing of TOPGAP significantly reduces cell death induced by TIKI. (A) Representative images of *N. benthamiana* leaves infiltrated with TIKI or EV control, along with RNAi:TOPGAP or the silencing control RNAi:GUS. Daylight and UV images were captured at 3 dpi, and cell death was quantified. (B) Box-and-dot plot showing that silencing TOPGAP significantly reduces cell death triggered by TIKI (0.29, n = 26), compared to the GUS-silencing control (9.65, n = 26). EV controls do not induce visible cell death. (C–D) Overexpression of TOPGAP increases HR induced by TIKI, dependent on its GAP activity. (C) Representative images of *N. benthamiana* leaves infiltrated with TIKI, with TOPGAP, TOPGAP^GAP^, or EV. Daylight and UV images were captured at 3 dpi, and HR was quantified. (D) Box-and-dot plot showing that overexpression of TOPGAP significantly increases HR triggered by TIKI (6.62, n = 22), compared to the EV control (3.30, n = 22), while the GAP mutant significantly reduces HR triggered by TIKI (0.607, n = 22). (E–F) Overexpression of TOPGAP increases HR induced by TIKI dependent on their interaction. (E) Representative images of *N. benthamiana* leaves infiltrated with TIKI, with TOPGAP, TOPGAP^7M^, or EV. Daylight and UV images were captured at 3 dpi, and HR was quantified. (F) Box-and-dot plot showing that overexpression of TOPGAP significantly increases HR triggered by TIKI (4.46, n = 18), compared to the EV control (1.68, n = 18), while the mutant TOPGAP^7M^ does not have any significant effect on the HR triggered by TIKI (1.34, n = 18). (G) AF2-M–predicted model of TIKI (green) targeting the RBD of TOPGAP (pink), depicting the interacting residues on TIKI. The bonds between TIKI and TOPGAP are predicted by PyMOL with a distance of 3 Å. TIKI interacts with TOPGAP using six key residues on TIKI, colored as cyan. (H) Substitution of six predicted interface residues strongly reduces association of TIKI with TOPGAP. GFP:TOPGAP was transiently co-expressed with either RFP:TIKI, RFP:TIKI^6A^, or RFP:EV. (I–J) The interaction-deficient mutant TIKI^6A^ does not cause visible cell death. (I) Representative images of *N. benthamiana* leaves infiltrated with TIKI, TIKI^6A^ or EV control. Daylight and UV images were captured at 3 dpi, and cell death was quantified. (J) Box-and-dot plot showing that while TIKI causes cell death (7.41, n = 21), TIKI^6A^ and the EV control do not cause visible cell death. (K–L) Overexpression of NRC4 enhances TIKI-triggered HR in WT plants. (K) Representative images of WT *N. benthamiana* leaves infiltrated with TIKI or EV control, together with NRC4 or EV. Daylight and UV images were captured at 3 dpi and HR was quantified. (L) Box-and-dot plot showing that NRC4 overexpression significantly increases HR triggered by TIKI (6.01, n = 19) compared with EV control (4.05, n = 19). (M–N) Genetic loss and complementation establish a contribution of NRC4 to TIKI-triggered HR. (M) Representative images of *nrc4* KO plants infiltrated with TIKI or EV, together with NRC4 or EV. In *nrc4* KO plants, TIKI-induced HR is reduced compared with WT, and reintroduction of NRC4 restores robust HR. Images were captured at 3 dpi and HR was quantified. (N) Box-and-dot plot showing that TIKI-induced HR is significantly attenuated in *nrc4* KO plants expressing EV (0.698, n = 27), whereas complementation with NRC4 restores and significantly enhances HR (7.58, n = 27). Co-expression of AVRblb2 and Rpi-blb2 was used as an HR control. (O–P) NRC2 and NRC3 do not substitute for NRC4 in supporting TIKI-triggered HR. (O) Representative images of *nrc2/3/4* KO plants infiltrated with TIKI together with NRC2, NRC3, NRC4 or EV. Images were captured at 3 dpi, and HR was quantified. (P) Box-and-dot plot showing that only NRC4 expression significantly enhances TIKI-triggered HR (6.43, n = 13) compared with EV (2.55, n = 13), whereas NRC2 (2.97, n = 13) and NRC3 (3.30, n = 13) do not significantly alter the response. Co-expression of PVX-CP and Rx was used as an HR control. Each color represents an independent biological replicate. Statistical comparisons were performed using Mann– Whitney U tests with Bonferroni correction in R. **P < 0.01; ***P < 0.001. ns, not significant.

We next tested the contribution of effector–target association. Unlike wild-type TOPGAP, the interaction-deficient mutant TOPGAP^7M^ did not significantly enhance TIKI-triggered cell death compared with EV (Figs. 4E, F), indicating that TOPGAP-mediated enhancement requires an interaction-competent host protein.

To examine this requirement from the effector side, we used AlphaFold-Multimer to guide mutagenesis of the predicted TIKI–TOPGAP interface. The model identified six candidate interface residues in TIKI, which we substituted with alanine to generate TIKI^6A^ (Fig. 4G; Fig. S10). Wild-type TIKI co-immunoprecipitated with TOPGAP, whereas TIKI^6A^ showed strongly reduced association (Fig. 4H). This reduction was accompanied by loss of detectable cell-death activity: TIKI^6A^ did not elicit visible cell death while wild-type TIKI induced a strong response (Figs. 4I, J). Together, these complementary perturbations of the host protein and effector support a requirement for TOPGAP engagement in TIKI-triggered cell death.

Having established the contribution of TOPGAP, we asked whether TIKI-triggered cell death also engages NRC4. Co-expression of NRC4 significantly enhanced the response in wild-type plants, whereas NRC4 did not induce detectable cell death when expressed with EV (Figs. 4K, L). In *nrc4* KO plants, TIKI retained limited cell-death activity, but reintroduction of NRC4 significantly enhanced the response, producing robust tissue collapse (Figs. 4M, N). These results identify NRC4 as a major contributor to TIKI-triggered cell death.

To determine whether other NRC helpers could substitute for NRC4, we co-expressed TIKI with NRC2, NRC3 or NRC4 in *nrc2/3/4* KO plants. Only NRC4 significantly enhanced TIKI-triggered cell death relative to EV; NRC2 and NRC3 did not significantly alter the response (Figs. 4O, P). Thus, as observed for PiE355, NRC2 and NRC3 did not substitute for NRC4 under the conditions tested.

We then tested whether robust TIKI-triggered cell death depends on the MADA-mediated execution function of NRC4 using the signaling-impaired mutant NRC4^L9E^. In *nrc4* KO plants, wild-type NRC4 supported strong TIKI-triggered cell death, whereas the response supported by NRC4^L9E^ was significantly weaker (Fig. S11). Neither NRC4 variant induced detectable cell death when co-expressed with EV. These findings connect TIKI-triggered cell death to the MADA-dependent execution activity of NRC4.

Together, these results extend the TOPGAP–NRC4 immune signaling module to a sequence-divergent effector homolog from another *Phytophthora* species. The shared contributions of TOPGAP catalytic activity, effector association and NRC4 execution support a model in which targeting of TOPGAP by related pathogen effectors engages a common immune surveillance mechanism.

## Discussion

We previously showed that PiE354 exploits the GAP activity of the host RabGAP TOPGAP to divert Rab8a-dependent defense secretion away from the pathogen interface^10^. Here we show that two related effectors, PiE355 from *P. infestans* and TIKI from *P. palmivora*, trigger cell death that depends on TOPGAP and its GAP activity. The C-terminal TBC domain of TOPGAP, which carries the GAP activity, also triggers cell death when expressed alone, and the N-terminal Rab-binding domain (RBD) that the effectors bind restrains this response. The helper NLR NRC4 mediates these responses, at least in part. We propose that binding of PiE355 and TIKI to the RBD alters the regulation of TOPGAP, thereby promoting immune activation (Fig. 5). PiE354 also binds the RBD and exploits GAP activity to promote susceptibility^10^, but does not elicit detectable cell death. The basis of this difference remains unresolved. Together, these findings link an effector-exploited trafficking susceptibility factor to NRC4-mediated immune signaling and support a model of surveillance of deregulated TOPGAP activity.

**Figure 5.**
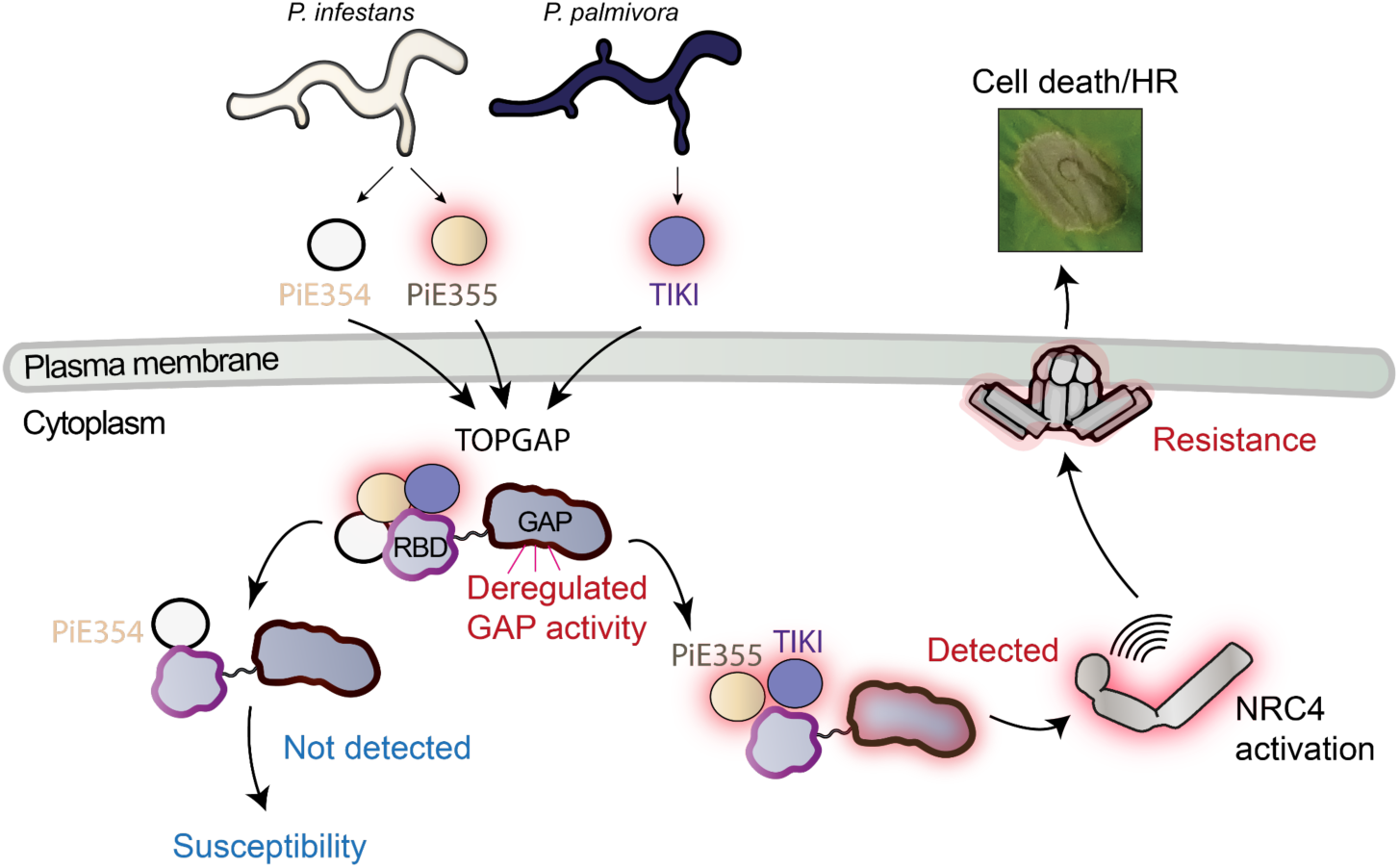
Proposed model for NRC4-mediated surveillance of TOPGAP deregulation by *Phytophthora* effectors. The *P. infestans* effectors PiE354 and PiE355 and the *P. palmivora* effector TIKI associate with the N-terminal Rab-binding domain (RBD) of TOPGAP. We propose that effector binding alters regulation of the C-terminal TBC (GAP) domain. PiE355 and TIKI elicit cell death involving TOPGAP catalytic activity and the helper NLR NRC4. PiE354 exploits TOPGAP to promote susceptibility but does not elicit detectable cell death. The mechanism underlying these contrasting outcomes remains unresolved, and an unidentified sensor NLR may act upstream of NRC4. Red shading indicates proposed activated states.

### What does the plant monitor in TOPGAP?

Silencing TOPGAP reduced the cell death triggered by PiE355 and TIKI, whereas overexpression of wild-type TOPGAP enhanced the response (Figs. 1A–D; Figs. 4A–D). This enhancement required GAP activity and effector binding, as the GAP-inactive mutant suppressed cell death and the interaction-deficient mutant TOPGAP^7M^ failed to enhance cell death (Figs. 1C–F; Figs. 4C–F). Mutating the predicted interface residues of the effectors had a matching effect: PiE355^7A^ lost detectable association with TOPGAP, TIKI^6A^ showed strongly reduced association, and neither triggered cell death (Figs. 1G–J; Figs. 4G–J). Neither TOPGAP nor GAP-inactive TOPGAP significantly altered the cell death triggered by AVRblb2–Rpi-blb2 or RxLR24 (Fig. S2), indicating that the enhancement does not extend to the other effector-triggered responses tested.

The TBC domain expressed alone triggered cell death without any effector, whereas full-length TOPGAP did not (Fig. 3B; Fig. S9)^10^. Substituting the catalytic residues R446 and Q483 abolished this response despite stable expression of the mutant (Figs. 3B, C; Fig. S6), and supplying the RBD *in trans* reduced the response without lowering the level of the TBC domain (Figs. 3D, E; Fig. S7). These findings indicate that the RBD restrains the cell death elicited by a catalytically active TBC domain, in line with the regulatory role suggested for the RBD in our earlier work^10^, supporting a link between regulation of the catalytic domain and immune activation. Further studies are needed to determine whether the RBD acts by limiting the access of Rab8a to the catalytic interface or by direct inhibition, and whether the plant senses TOPGAP itself or the resulting change in Rab-dependent trafficking.

### How is TOPGAP activity relayed to NRC4?

NRC4 is the principal NRC helper for PiE355- and TIKI-triggered cell death. Reintroducing NRC4 into *nrc4* KO plants restored robust responses to both effectors, whereas NRC2 and NRC3 did not substitute in *nrc2/3/4* KO plants (Figs. 2C–F and Figs. 4M–P). The MADA-motif mutant NRC4^L9E^, which is impaired in cell death execution^22^, restored significantly weaker responses than wild-type NRC4, consistent with signaling through the NRC4 resistosome (Figs. S4 and S11). NRC4 co-expression also enhanced the cell death triggered by the TBC domain without increasing TBC domain abundance (Figs. 3F, G and Fig. S8), and this response was weaker in *nrc4* KO plants (Fig. S9). Loss of NRC4 weakened but did not abolish the cell death triggered by PiE355, TIKI and the TBC domain, indicating that NRC4 mediates each response at least in part.

How TOPGAP activity reaches NRC4 is not known. NRC helpers generally act downstream of sensor NLRs^20,33^. Although IP–MS identified NRC4 as a candidate TOPGAP interactor, this does not show direct binding or recognition, and an unidentified sensor NLR could detect a change in TOPGAP or its downstream effects and signal through NRC4.

### Why do TOPGAP-targeting effectors differ in their ability to trigger cell death?

TIKI and PiE355 share only about 50% amino acid identity across their effector domains, yet both engage the TOPGAP–NRC4 signaling module. TOPGAP enhances cell death triggered by either effector, and this enhancement requires both its GAP activity and its ability to bind the effectors. Loss of NRC4 reduces these cell death triggered by both effectors whereas its overexpression boosts it, indicating that the TOPGAP–NRC4 module responds to effectors from two *Phytophthora* species. The shared requirement for effector binding and GAP activity points to the catalytic state of effector-bound TOPGAP as a potential immune cue, linking sequence-divergent effectors to a common NRC4-mediated response.

Intriguingly, PiE354 also binds TOPGAP but does not trigger detectable cell death. It associates with the RBD and exploits TOPGAP to promote susceptibility^10^, yet elicits no detectable cell death, even when TOPGAP or NRC4 is co-expressed (Fig. S1; Figs. 2A and 2B). Because the three effectors share a common fold and a binding interface^10^, one possibility is that they differ in the extent of TOPGAP deregulation. Our earlier work proposed that PiE354 docking at the RBD may relieve restraint by this domain and guide Rab8a to the catalytic interface^10^. A more limited or spatially restricted change in GAP activity could promote susceptibility without triggering cell death. Such differences in catalytic regulation could be coupled to distinct conformations of effector-bound TOPGAP, some of which may be recognized by a sensor NLR. Direct comparisons of how the three effectors affect TOPGAP activity and conformation are needed to establish why PiE355 and TIKI trigger cell death whereas PiE354 does not.

### How does TOPGAP surveillance relate to established guard mechanisms?

Our model places TOPGAP within the guard framework, in which NLRs monitor host proteins that effectors target^36^. In plants, ZAR1 is activated when the effector AvrAC uridylylates the decoy kinase PBL2, which is then recruited to the ZAR1–RKS1 complex^21^. In animals, the Pyrin inflammasome responds to the inactivation of RhoA by bacterial toxins and so reports the state of a host GTPase rather than a microbial molecule^37^. In both examples the cue is a modification of a host protein or a loss of its activity. In the proposed TOPGAP model, the host enzyme retains the catalytic activity exploited during infection, and our data implicate deregulation of that activity in immune activation. Whether the surveillance machinery senses TOPGAP conformation, catalytic output or downstream trafficking changes remains to be established.

Our findings connect the catalytic function of an effector-exploited trafficking susceptibility factor to NLR-mediated immune signaling. PiE355 and TIKI engage a shared TOPGAP–NRC4 module despite sequence divergence, whereas PiE354 associates with TOPGAP without eliciting detectable cell death. These observations support a model in which perturbation of host enzyme regulation can provide a cue for immune surveillance. Other trafficking regulators whose catalytic activities are exploited by effectors may be monitored in a similar way. Defining how effectors alter TOPGAP function and how these changes engage NRC4 will be essential for testing this model. This knowledge could inform strategies to reduce effector exploitation of TOPGAP or strengthen immune responses to its manipulation while preserving its normal trafficking functions.

## Materials and Methods

### Plant growth details

*Nicotiana benthamiana* plants (wild-type and transgenic lines) were grown in a controlled growth chamber maintained at 24 °C. Plants were cultivated in a soil mixture consisting of organic compost supplemented with sand and vermiculite (Levington’s F2 with sand and Sinclair’s 2–5 mm vermiculite at a 3:1 ratio). Plants were grown under high-intensity light with a long-day photoperiod of 16 h light and 8 h dark. All experiments were performed using 4-week-old plants. Transgenic lines used in this study included *nrc4* CRISPR knockout plants^22^, and *nrc2/3/4* CRISPR knockout plants^38^.

### Molecular cloning

TBCDF^GAP^, PiE355^7A^, TIKI^6A^ and RxLR24 constructs were generated using Gibson Assembly, following previously established protocols^39^. A pK7WGF2-derived expression vector, domesticated for Gibson Assembly and encoding an N-terminal fluorescent tag, was used as the cloning backbone. Target sequences were synthesized as gene fragments (Table S2) and assembled into the vector using Gibson Assembly. The resulting constructs were introduced into *Escherichia coli* DH5α chemically competent cells by heat shock. Plasmids were amplified and purified using the PureYield™ Plasmid Miniprep System (Promega), then transferred into *Agrobacterium tumefaciens* strain GV3101 by electroporation. Construct sequences were verified by Sanger sequencing (Eurofins) or full-length plasmid sequencing (Full Circle Labs). *Agrobacterium* transformants were selected on LB agar supplemented with gentamicin and spectinomycin. A complete list of constructs used in this study is provided in Table S3.

### Agrobacterium-mediated transient gene expression in *N. benthamiana*

Transient expression assays were performed in *N. benthamiana* by *Agrobacterium*-mediated leaf infiltration as described previously^40^. *Agrobacterium tumefaciens* strains carrying the indicated constructs were pelleted, washed with water, and resuspended in infiltration buffer containing 10 mM MES and 10 mM MgCl₂ (pH 5.7). Bacterial density was measured using a BioPhotometer spectrophotometer (Eppendorf) and adjusted to the appropriate optical density (OD₆₀₀) for each construct and experimental setup. The suspensions were infiltrated into fully expanded leaves of 4-week-old plants using a needleless 1-ml syringe.

### Co-immunoprecipitation and immunoblot analyses

Proteins were transiently expressed in *N. benthamiana* leaves by agroinfiltration and harvested at 3 days post infiltration (dpi). For co-immunoprecipitation assays, 500 mg of leaf tissue was collected per sample. Protein extraction, immunoprecipitation, and immunoblot analyses were performed as previously described^31^. Primary antibodies included mouse monoclonal anti-RFP and rabbit polyclonal anti-GFP antibodies (Chromotek). Secondary antibodies consisted of alkaline phosphatase (AP)-conjugated anti-rabbit and horseradish peroxidase (HRP)-conjugated anti-mouse antibodies (Sigma-Aldrich). A complete list of antibodies used in this study is provided in Table S4.

### Structural analyses

Structure predictions were generated using AlphaFold 3 (v3.0.2)^41^ and AlphaFold2-Multimer (AF2-M) in ColabFold v1.5.5^42^, accessed through a Google Colab subscription, following established AlphaFold guidelines^43^. Predicted structures were visualized in UCSF ChimeraX (version 1.12). Local model confidence was assessed using predicted local distance difference test (pLDDT) scores, which estimate the lDDT-Cα metric^44^. Scores range from 0 to 100, with higher values indicating greater model reliability. A complete list of protein sequences used for AlphaFold predictions is provided in Table S5.

### Cell death assay and data analysis

Cell death assays were conducted by agroinfiltrating the indicated constructs into the abaxial side of *N. benthamiana* leaves. At 2–4 dpi, infiltrated leaves were excised and imaged under daylight, ultraviolet (UV) and infrared conditions. Depending on the signal intensity of each experiment, hypersensitive response (HR) intensity was quantified from either the infrared or the UV images using ImageJ. Infrared signals were acquired using the 800-nm channel of a Bio-Rad imaging system, whereas UV signals were quantified by subtracting green fluorescence from red chlorophyll autofluorescence. Normalized HR intensities were plotted using ggplot2 in R. Group-wise normality was assessed by Shapiro-Wilk test and homogeneity of variance by Levene’s test. Because one or more groups in each dataset deviated from normality, pairwise Mann–Whitney *U* tests with Bonferroni correction for multiple comparisons were applied throughout. Statistical significance was indicated as \**P* < 0.05, \*\**P* < 0.01 and \*\*\**P* < 0.001. Details of all statistical calculations can be found in Table S6.

### Accession numbers

TOPGAP (Nbe.v1.s00100g29830), TIKI (PLTG_0964243), PiE354 (PITG_04354), PiE355 (PITG_04355), RxLR24 (XP002997548/PITG_18405).

## Acknowledgments

We thank members of the SchoBozKa community for insightful discussions and valuable suggestions throughout this work. We acknowledge the use of Claude (Anthropic) for proofreading the manuscript. E.L.H.Y. thanks Helena Wu for their support during manuscript preparation.

## Author contributions

Conceptualization: E.L.H.Y., T.O.B. Methodology: E.L.H.Y., N.E.

Validation: E.L.H.Y., Y.W. Formal Analysis: E.L.H.Y., N.E.

Investigation: E.L.H.Y., N.E., Y.W. Resources: E.L.H.Y.

Data Curation: E.L.H.Y. Visualization: E.L.H.Y.

Writing – Original Draft: E.L.H.Y., T.O.B.

Writing – Review & Editing: E.L.H.Y., N.E., T.O.B. Supervision: E.L.H.Y., T.O.B.

Funding Acquisition: E.L.H.Y., N.E., T.O.B.

## Data and materials availability

All relevant study data are included in the article and Supplementary Materials. Alphafold predictions are uploaded to the public repository figshare and available at https://doi.org/10.6084/m9.figshare.34113588.

## Supplementary Materials included in this manuscript

Supplementary Figures S1–S11. Supplementary Tables S2–S5.

## Additional Supplementary Materials supplied separately

Table S1. TOPGAP IP–MS dataset. Table S6. Summary of Statistics.

**Table S2.**
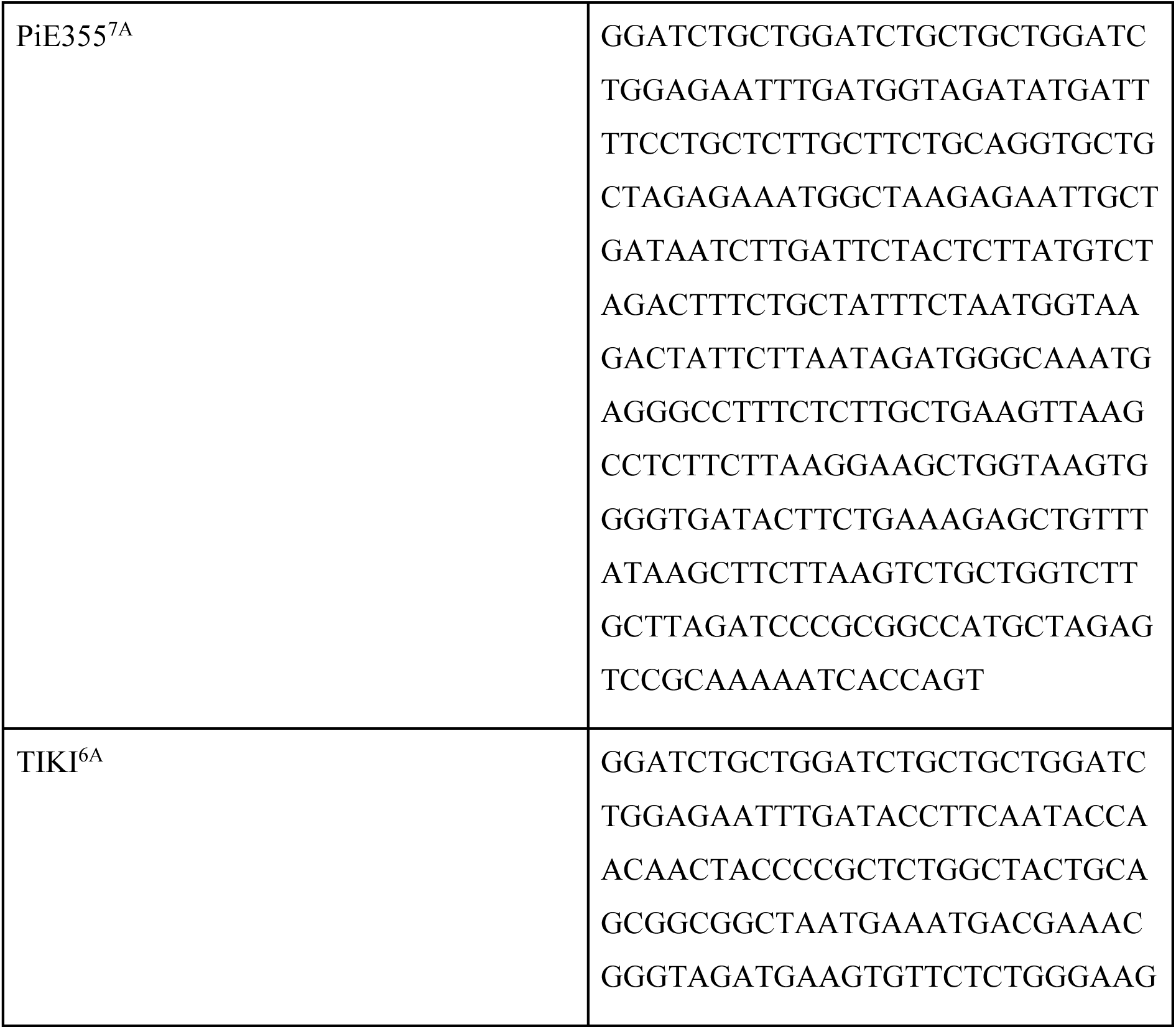
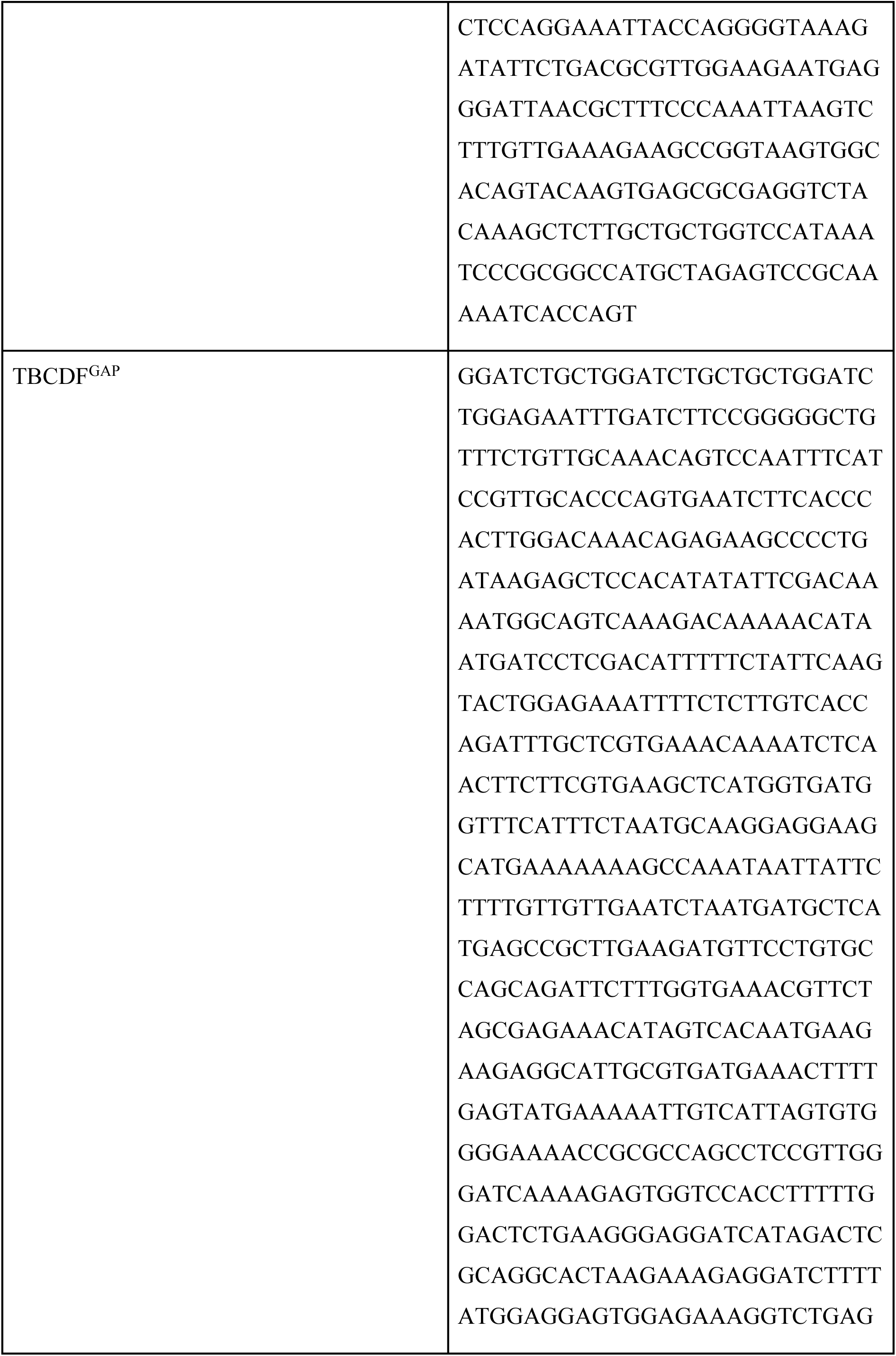
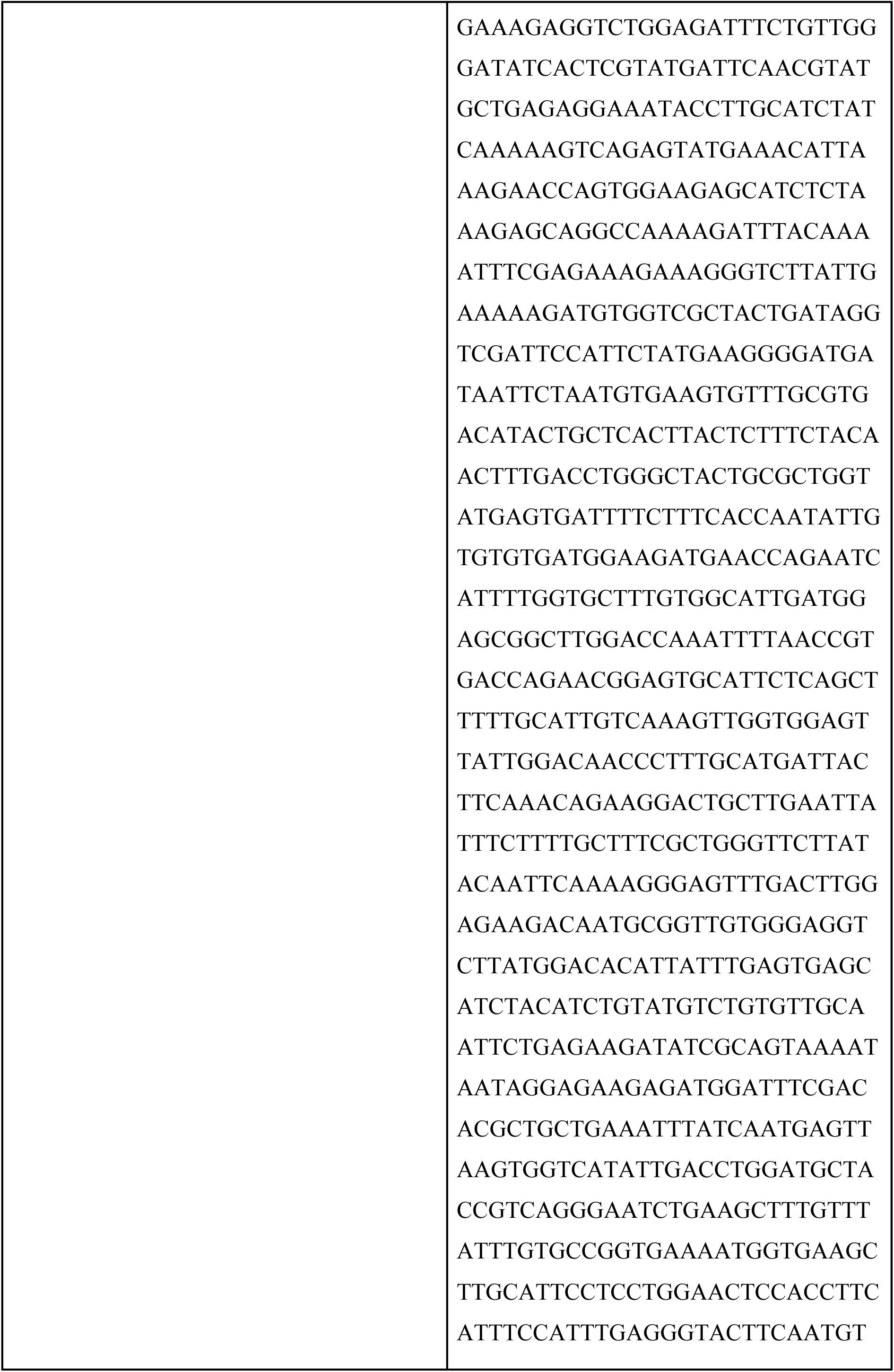
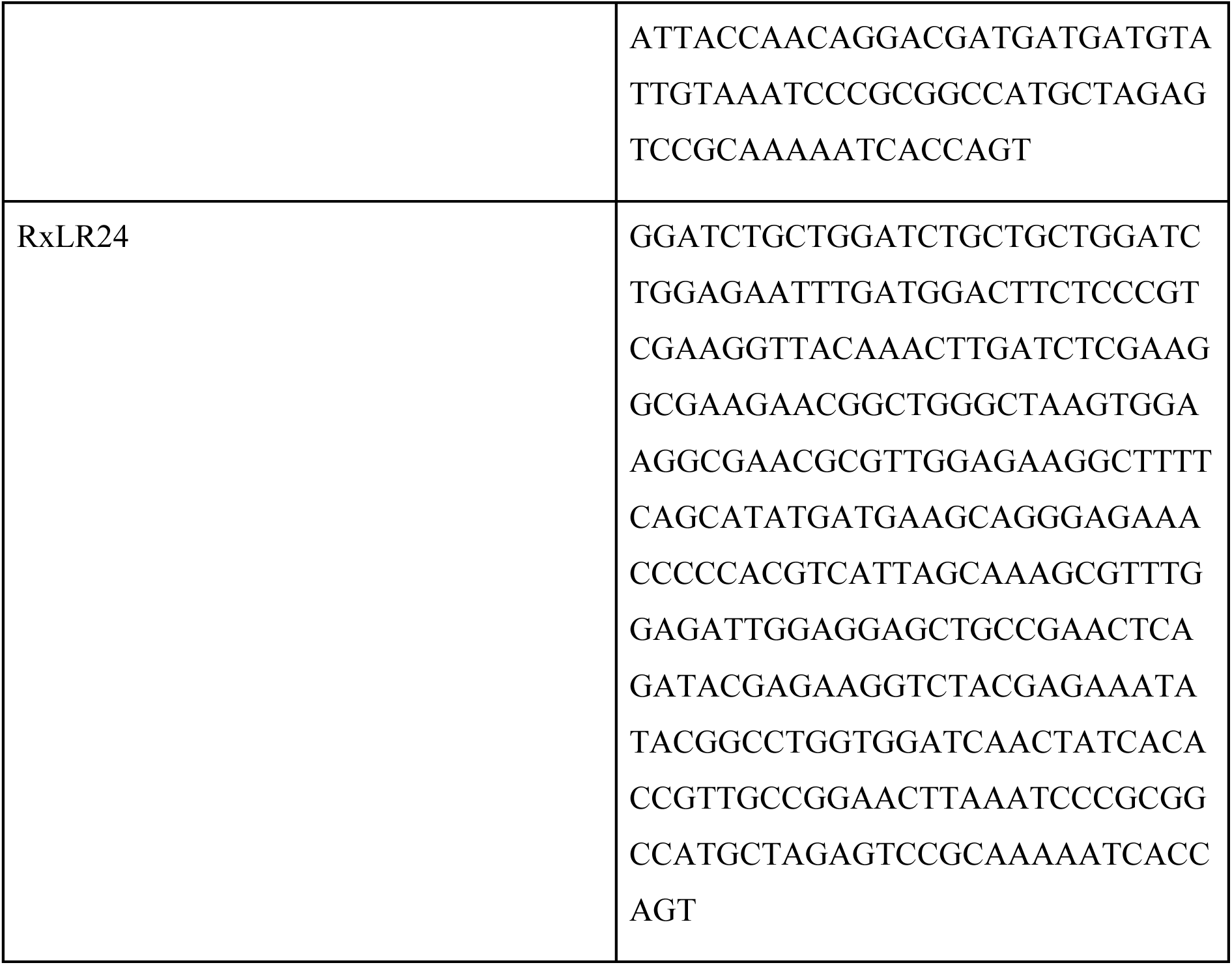
List of synthetic fragments used in this study.

**Table S3.** List of constructs used in this study.

|  |  |
| --- | --- |
| Vector (pK7WGF2 domesticated for Gibson assembly) | 45 |
| Vector (pRNAi-GG) | 46 |
| AVRblb2 | 47 |
| Rpi-blb2 | 47 |
| PVX-CP | 19 |
| Rx | 19 |
| RxLR24 | Cloned for this paper (synthetic fragment: RxLR24) |
| RNAi:TOPGAP | 10 |
| RNAi:GUS | 5 |
| RFP:PiE355 | 10 |
| RFP:PiE355<sup>7A</sup> | Cloned for this paper (synthetic fragment: PiE355<sup>7A</sup>) |
| RFP:PiE354 | 10 |
| RFP:TIKI | 10 |
| RFP:TIKI<sup>6A</sup> | Cloned for this paper (synthetic fragment: TIKI<sup>6A</sup>) |
| RFP:TOPGAP | 10 |
| RFP:TOPGAP<sup>GAP</sup> | 10 |
| RFP:TOPGAP<sup>7M</sup> | 10 |
| GFP:TOPGAP | 10 |
| NRC2 | 48 |
| NRC3 | 48 |
| NRC4 | 26 |
| NRC4<sup>L9E</sup> | 22 |
| GFP:RBDF | 10 |
| RFP:RBDF | 10 |
| RFP:TBCDF | 10 |
| RFP:TBCDF<sup>GAP</sup> | Cloned for this paper (synthetic fragment: TBCDF<sup>GAP</sup>) |
| GFP:EV | 49 |
| HA:EV | 49 |
| RFP:EV | 49 |

**Table S4.** List of antibodies used in this study.

| Primary/Secondary | Detail | Company | Product Code |
| --- | --- | --- | --- |
| Primary | Anti-RFP (Mouse monoclonal) | Chromotek | 6G6 |
| Primary | Anti-GFP (Rabbit polyclonal) | Chromotek | PABG1 |
| Secondary | Anti-rabbit (AP) | Sigma-Aldrich | SAB3700834 |
| Secondary | Anti-mouse (HRP) | Sigma-Aldrich | SAB3700986 |

**Table S5.**
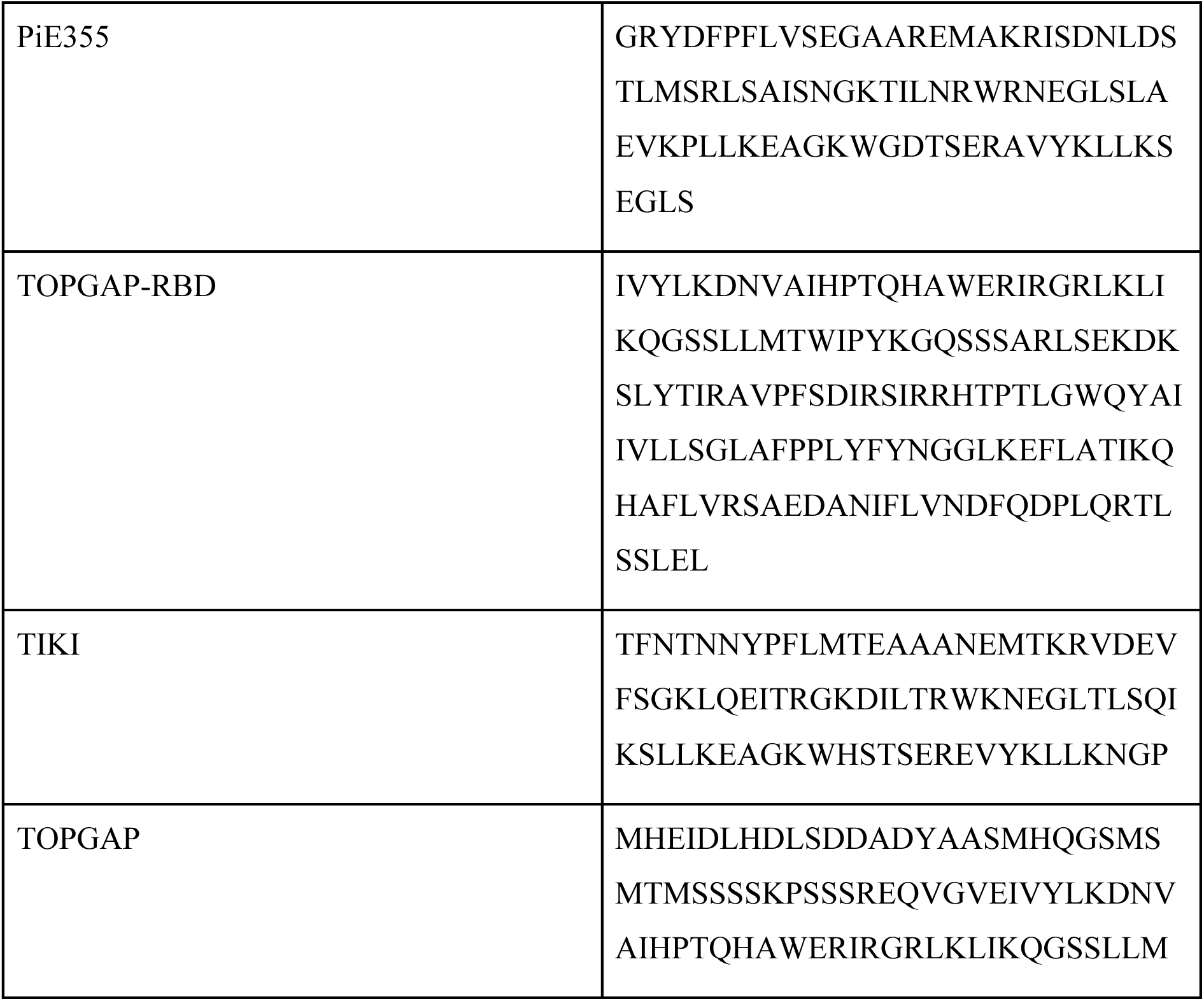
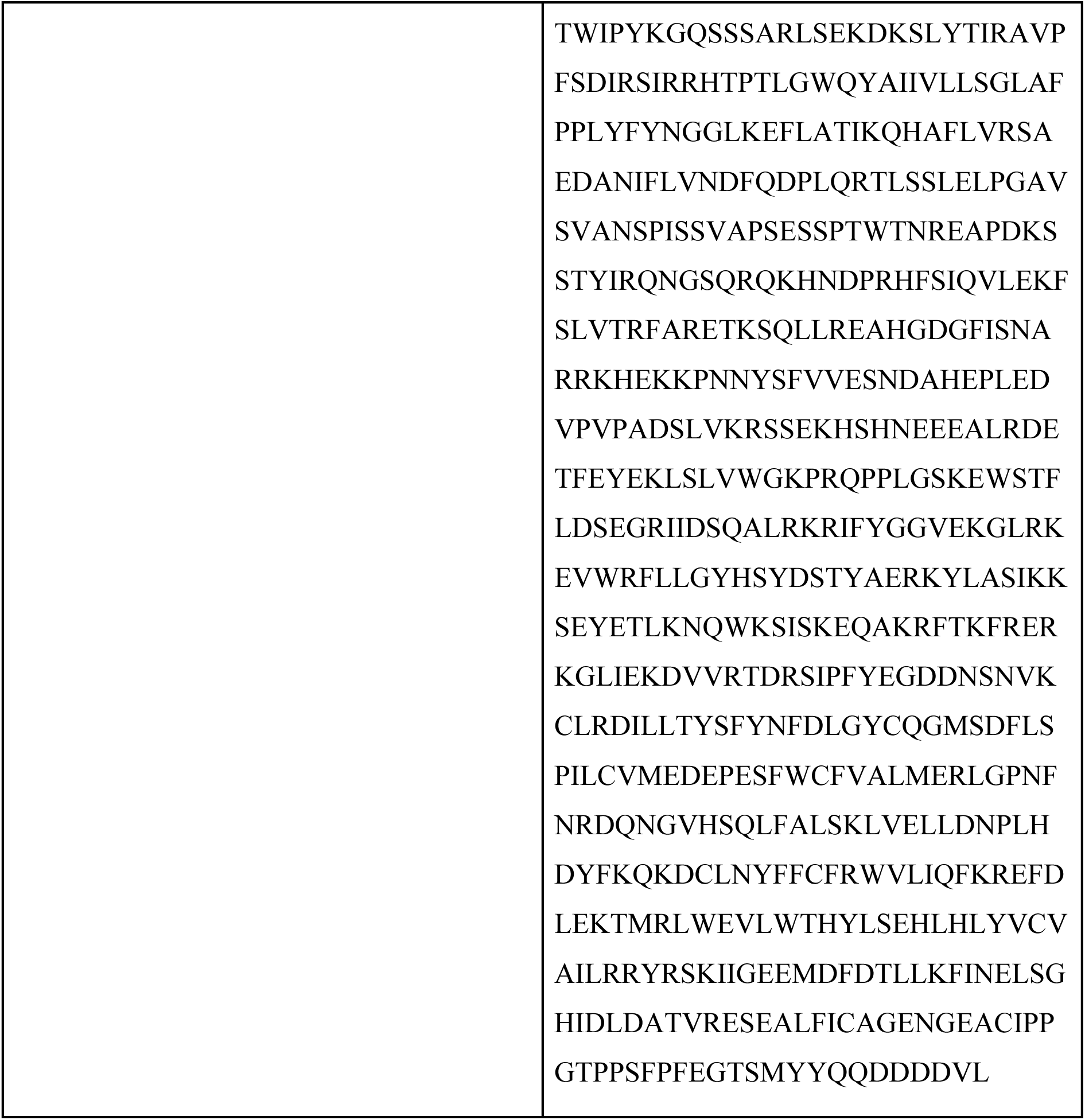
List of protein sequences used for AlphaFold predictions.

**Figure S1.**
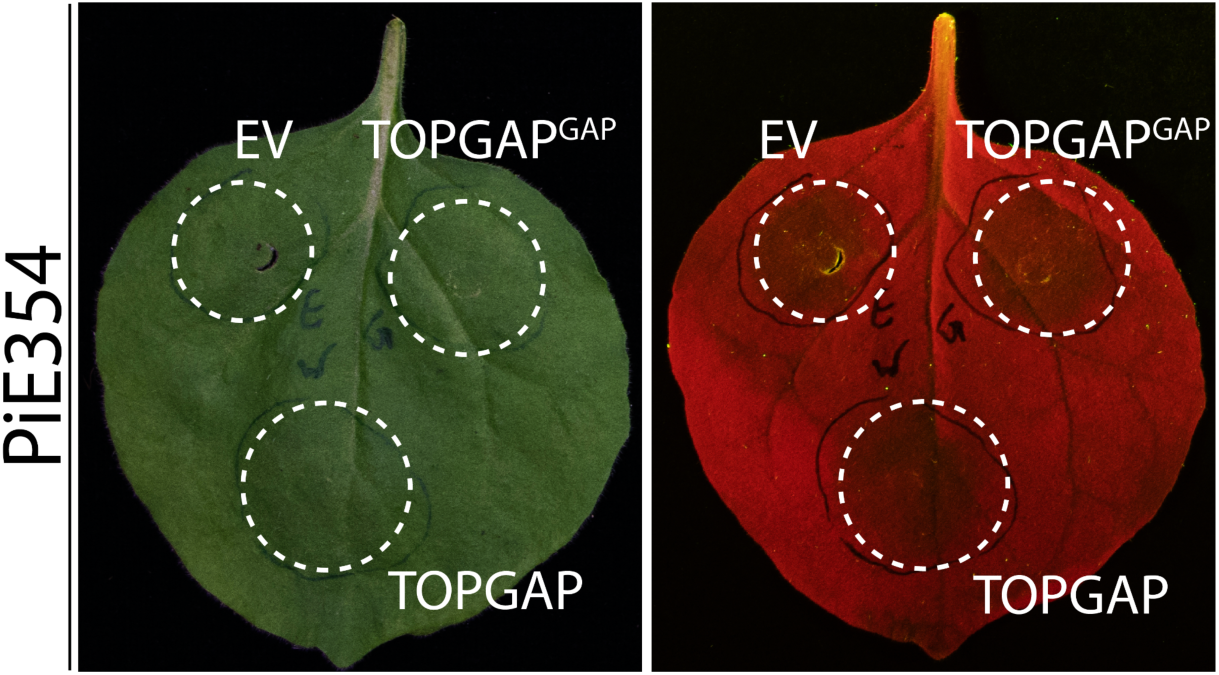
PiE354 does not elicit visible cell death upon co-expression with TOPGAP or its GAP-inactive mutant. Representative images of *N. benthamiana* leaves infiltrated with PiE354, with TOPGAP, TOPGAP^GAP^, or EV. Daylight and UV images were captured at 4 dpi. No visible cell death was observed in any infiltrated sample (n = 12).

**Figure S2.**
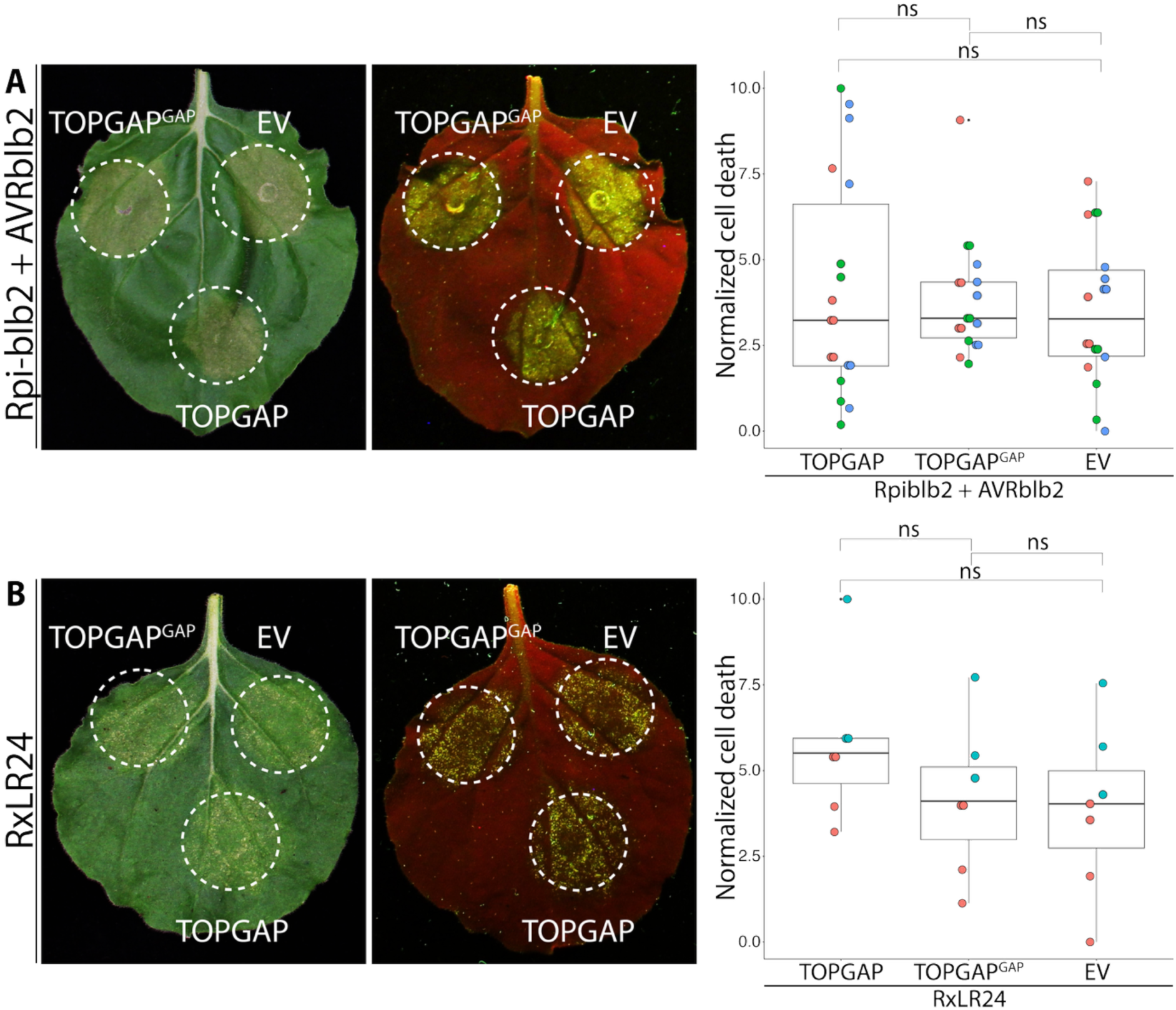
Overexpression of TOPGAP does not significantly alter cell death triggered by AVRblb2 or RxLR24. (A) Representative images of *N. benthamiana* leaves infiltrated with the sensor NLR Rpi-blb2 and its cognate effector AVRblb2, together with TOPGAP, the GAP-inactive mutant (TOPGAP^GAP^), or EV control. Daylight and UV images were captured at 4 dpi and HR was quantified. Box-and-dot plot showing that overexpression of TOPGAP (4.14, n = 18) or TOPGAP^GAP^ (3.85, n = 18) does not significantly alter HR triggered by AVRblb2 compared with EV control (3.52, n = 18). (B) Representative images of *N. benthamiana* leaves infiltrated with RxLR24, together with TOPGAP, TOPGAP^GAP^, or EV control. Daylight and UV images were captured at 4 dpi and HR was quantified. Box-and-dot plot showing that overexpression of TOPGAP (5.69, n = 7) or TOPGAP^GAP^ (4.16, n = 7) does not significantly alter HR triggered by RxLR24 compared with EV control (3.87, n = 7). Statistical comparisons were performed using Mann–Whitney U tests with Bonferroni correction in R. ns, not significant.

**Figure S3.**
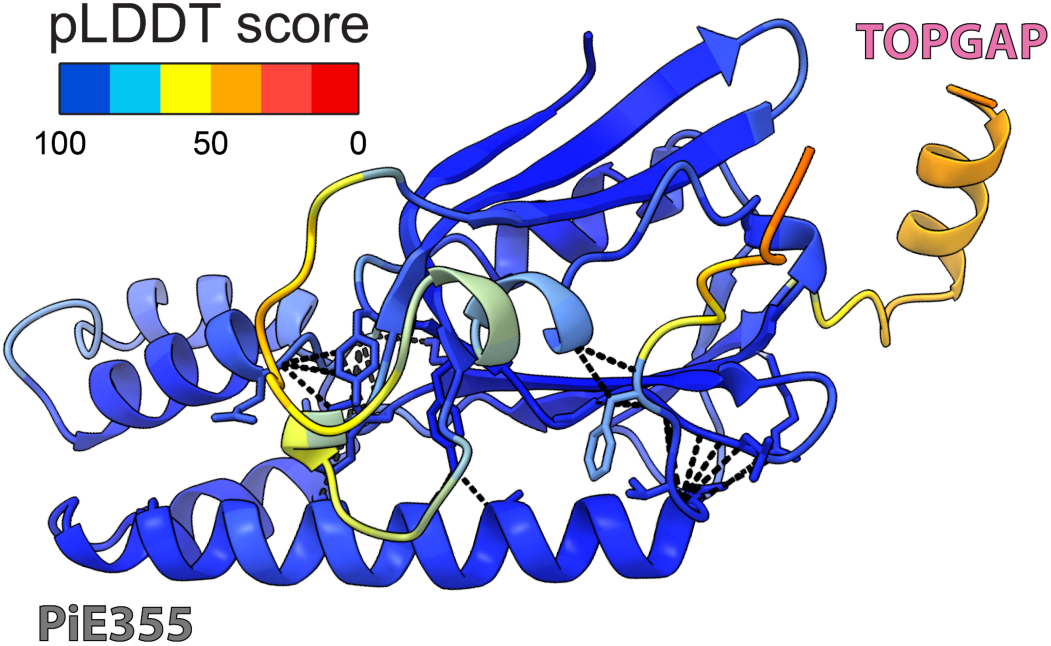
AF2-M–predicted model of PiE355 in complex with the RBD of TOPGAP. The PiE355–RBD structural model is colored according to the AlphaFold prediction confidence (pLDDT) scores, as indicated in the rectangular box.

**Figure S4.**
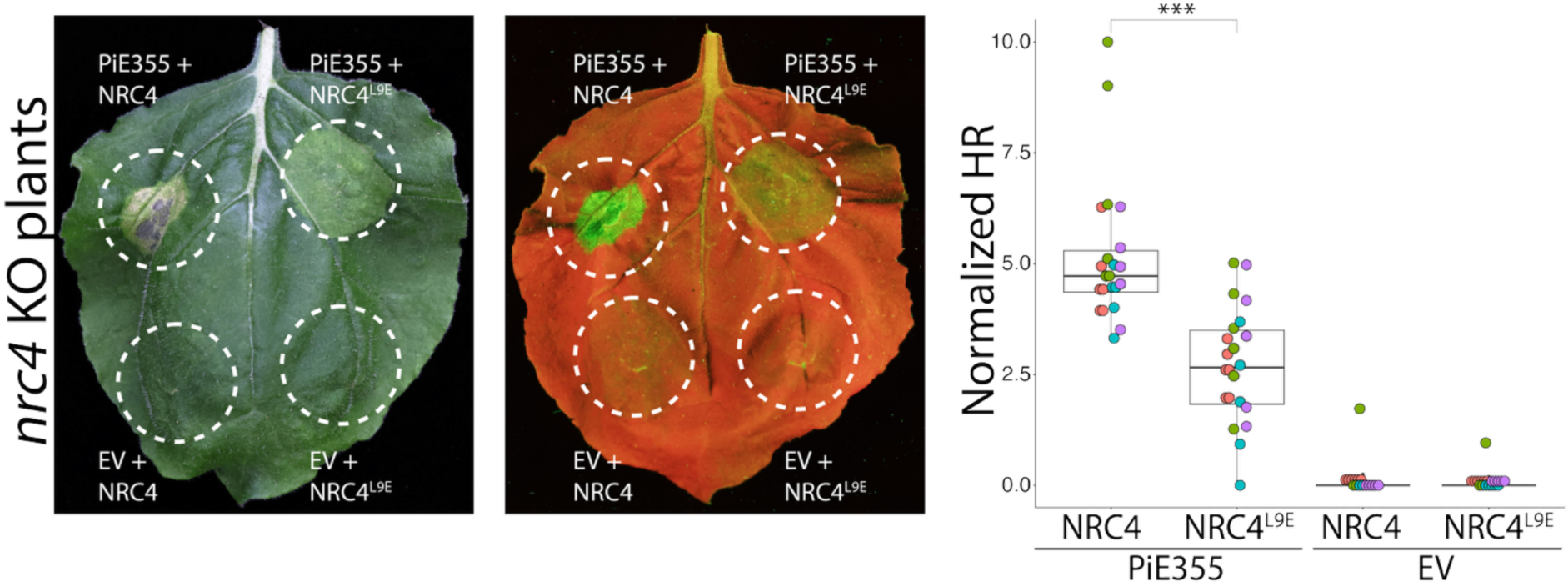
An intact MADA motif in NRC4 supports robust PiE355-triggered hypersensitive response. Representative images of *nrc4* KO *N. benthamiana* leaves infiltrated with PiE355 or EV control together with wild-type NRC4 or the MADA-motif mutant NRC4^L9E^. Daylight and UV images were captured at 4 dpi and HR was quantified. Reintroduction of wild-type NRC4 restores robust PiE355-triggered HR in *nrc4* KO plants, whereas expression of the NRC4^L9E^ mutant fails to restore robust HR, resulting in significantly reduced cell death. Box-and-dot plot showing that WT NRC4 significantly enhances PiE355-triggered HR (5.17, n = 22) compared with NRC4^L9E^ (2.72, n = 22). Expression of NRC4 or NRC4^L9E^ together with EV does not induce visible HR. Statistical comparisons were performed using Mann–Whitney U tests with Bonferroni correction in R. ***P < 0.001.

**Figure S5.**
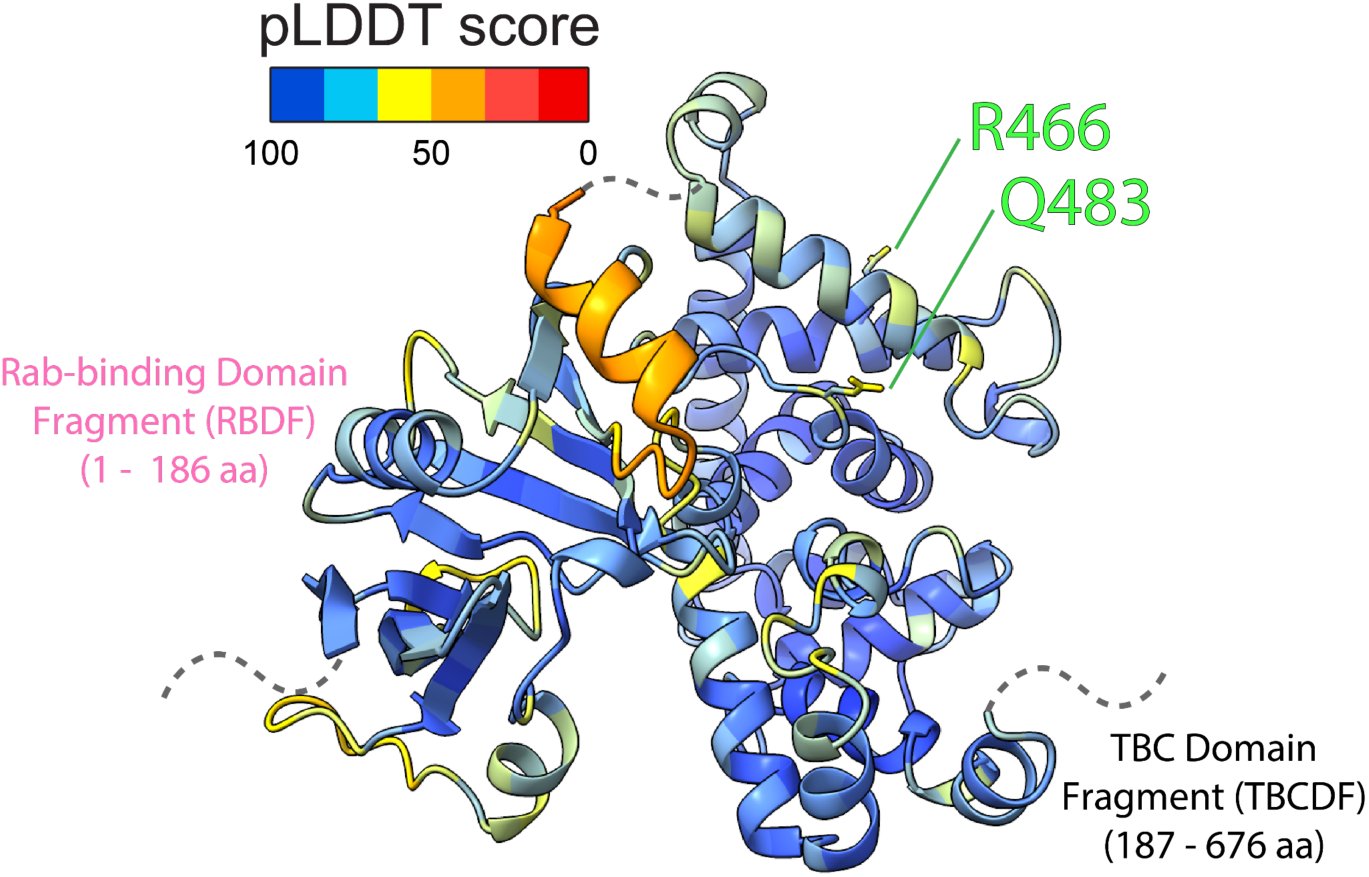
AF3–predicted model of TOPGAP. The TOPGAP structural model is colored according to the AlphaFold prediction confidence (pLDDT) scores, as indicated in the rectangular box.

**Figure S6.**
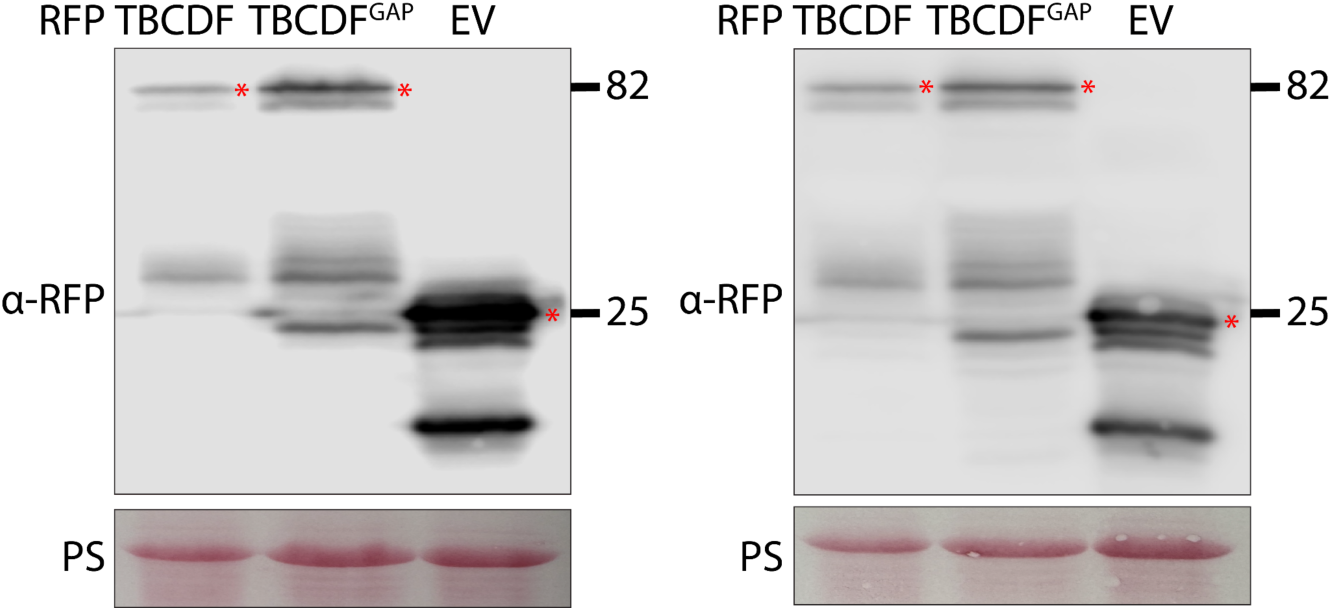
Western blot confirming robust expression of TBCDF^GAP^. RFP:TBCDF, RFP:TBCDF^GAP^, or RFP:EV was transiently expressed in plants. Total protein extracts were prepared at 3 dpi and immunoblotted. Red asterisks show expected band sizes. Numbers on the right indicate kDa values.

**Figure S7.**
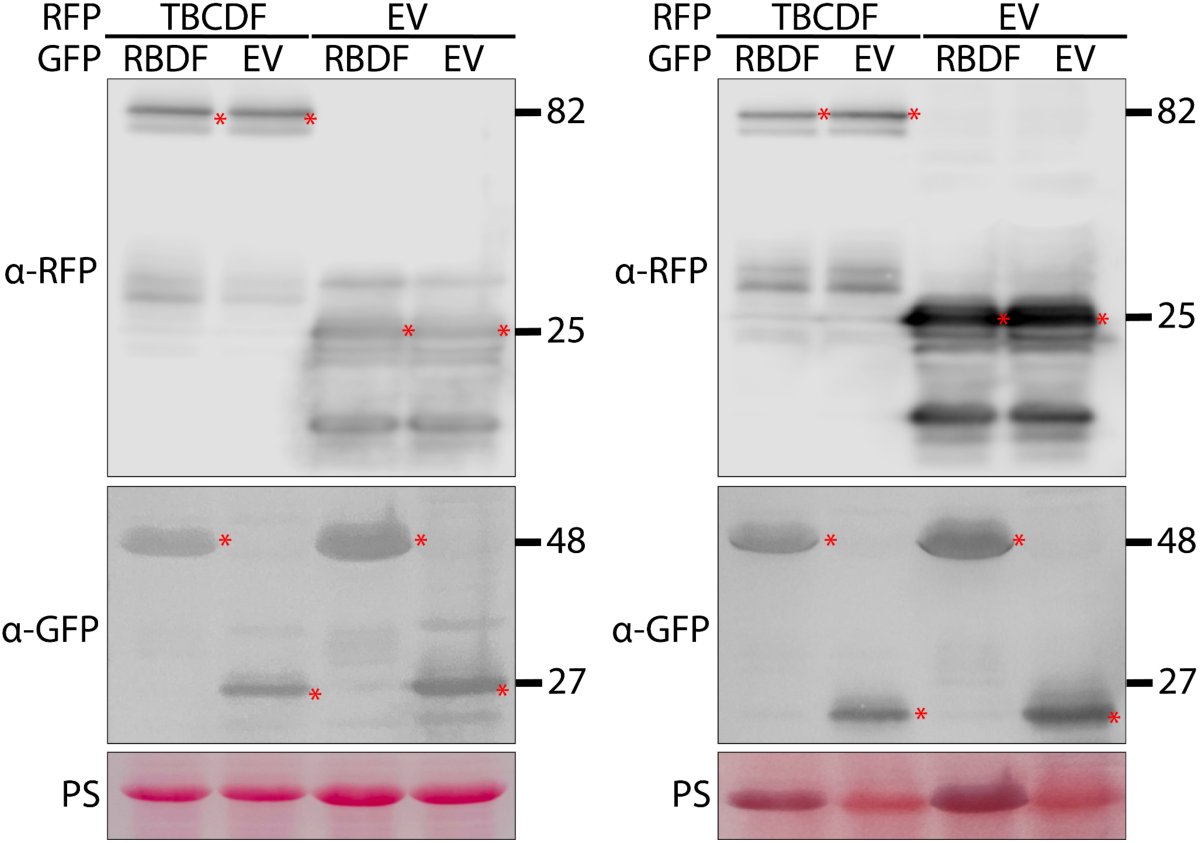
Western blot confirming comparable expression levels of TBCDF when co-expressed with RBDF or EV. RFP:TBCDF or RFP:EV was transiently co-expressed with GFP:RBDF or GFP:EV in plants. Total protein extracts were prepared at 3 dpi and immunoblotted. Red asterisks show expected band sizes. Numbers on the right indicate kDa values.

**Figure S8.**
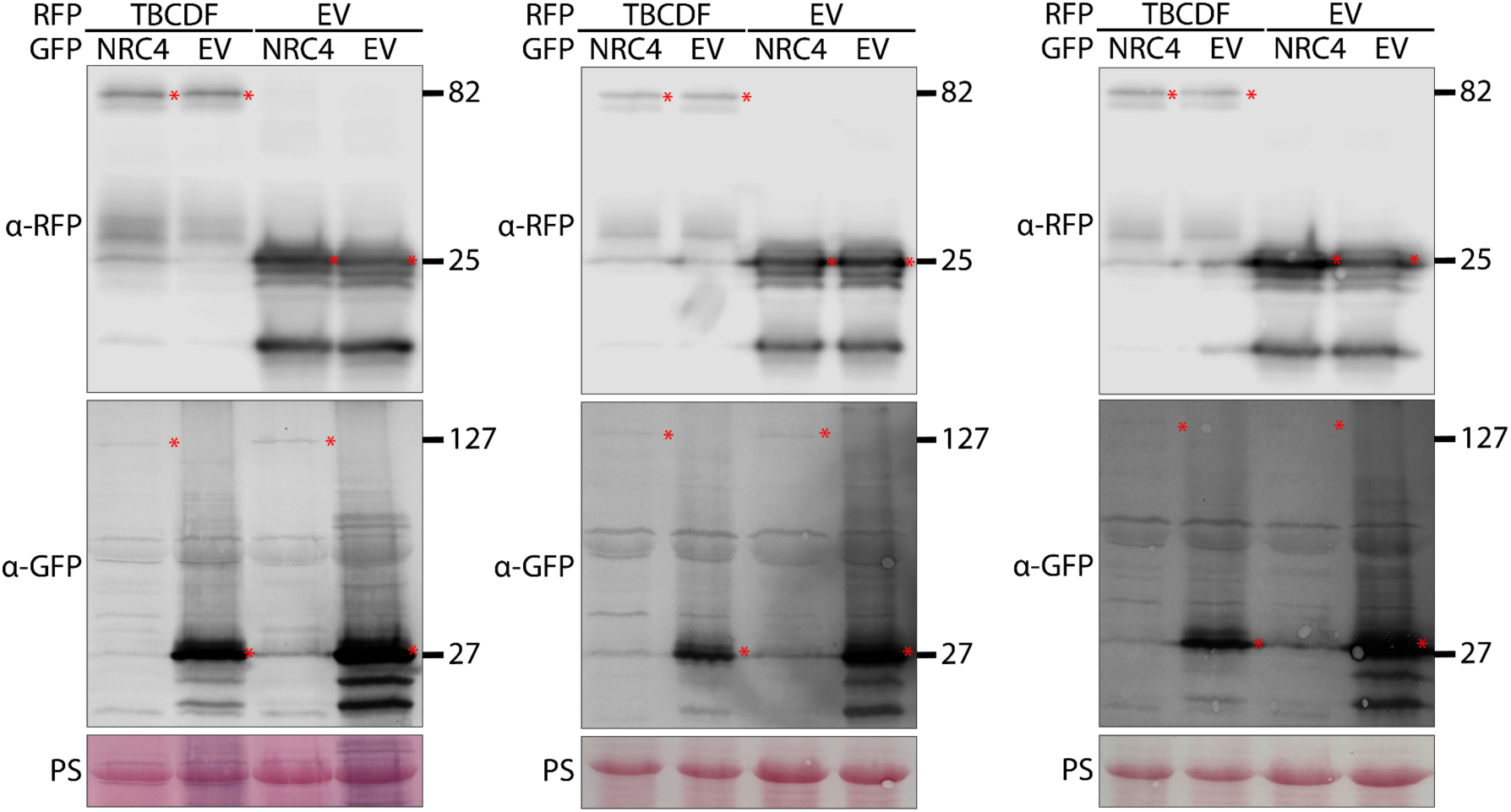
Western blot confirming comparable expression levels of TBCDF when co-expressed with NRC4 or EV. RFP:TBCDF or RFP:EV was transiently co-expressed with GFP:NRC4 or GFP:EV in plants. Total protein extracts were prepared at 3 dpi and immunoblotted. Red asterisks show expected band sizes. Numbers on the right indicate kDa values.

**Figure S9.**
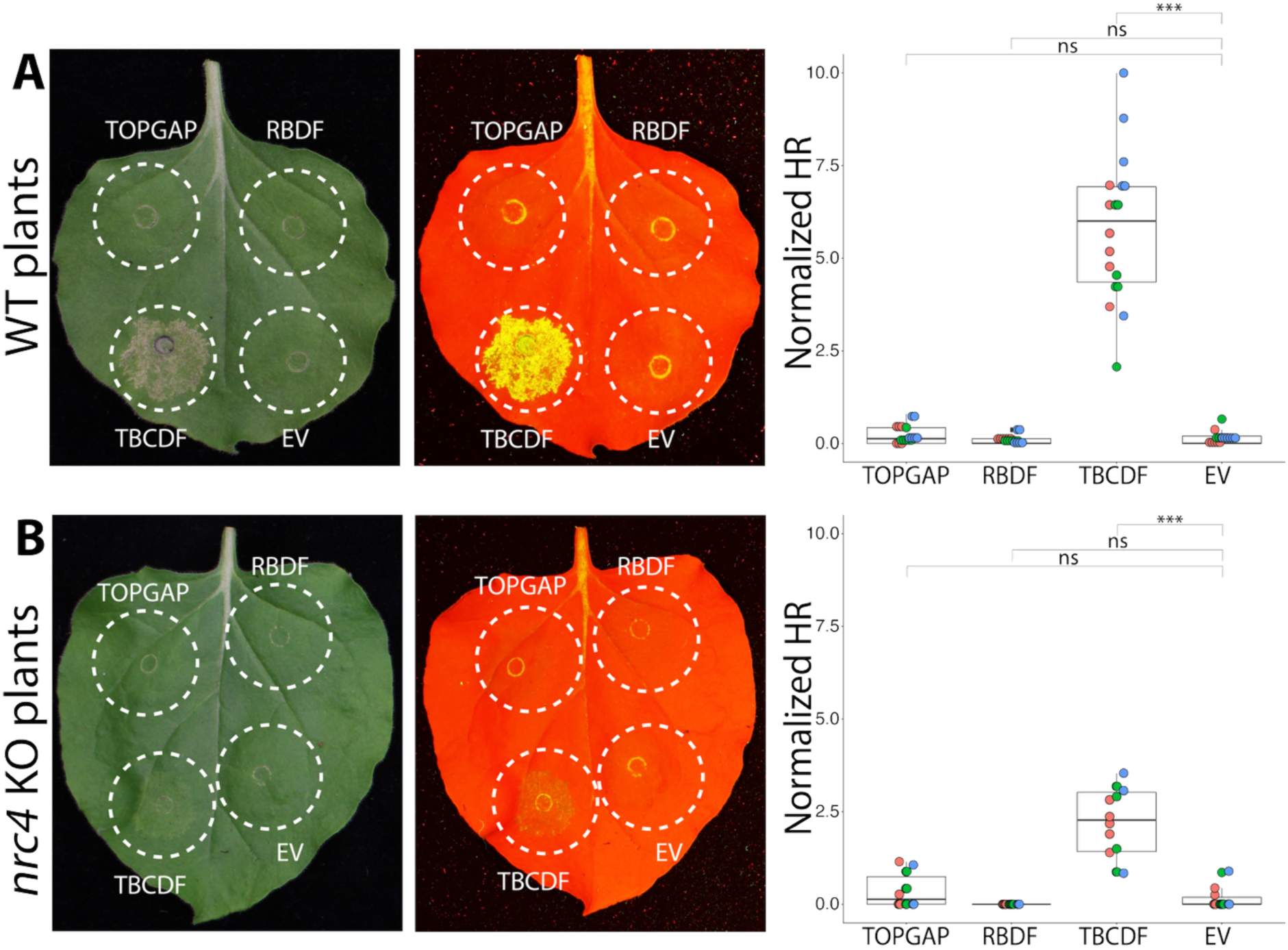
TBCDF-induced HR is attenuated in *nrc4* KO plants. Representative images of. (A) WT and (B) *nrc4* KO *N. benthamiana* leaves infiltrated with full-length TOPGAP, RBDF, TBCDF, or EV control. Images were captured at 4 dpi under daylight and UV illumination, and HR was quantified. Box-and-dot plot showing that TBCDF-induced HR is reduced in *nrc4* KO plants (2.19, n = 14) compared with WT plants (5.80, n = 18). Expression of full-length TOPGAP, RBDF, or EV does not induce visible cell death in either genotype. Statistical comparisons were performed using Mann–Whitney U tests with Bonferroni correction in R. ***P < 0.001. ns, not significant.

**Figure S10.**
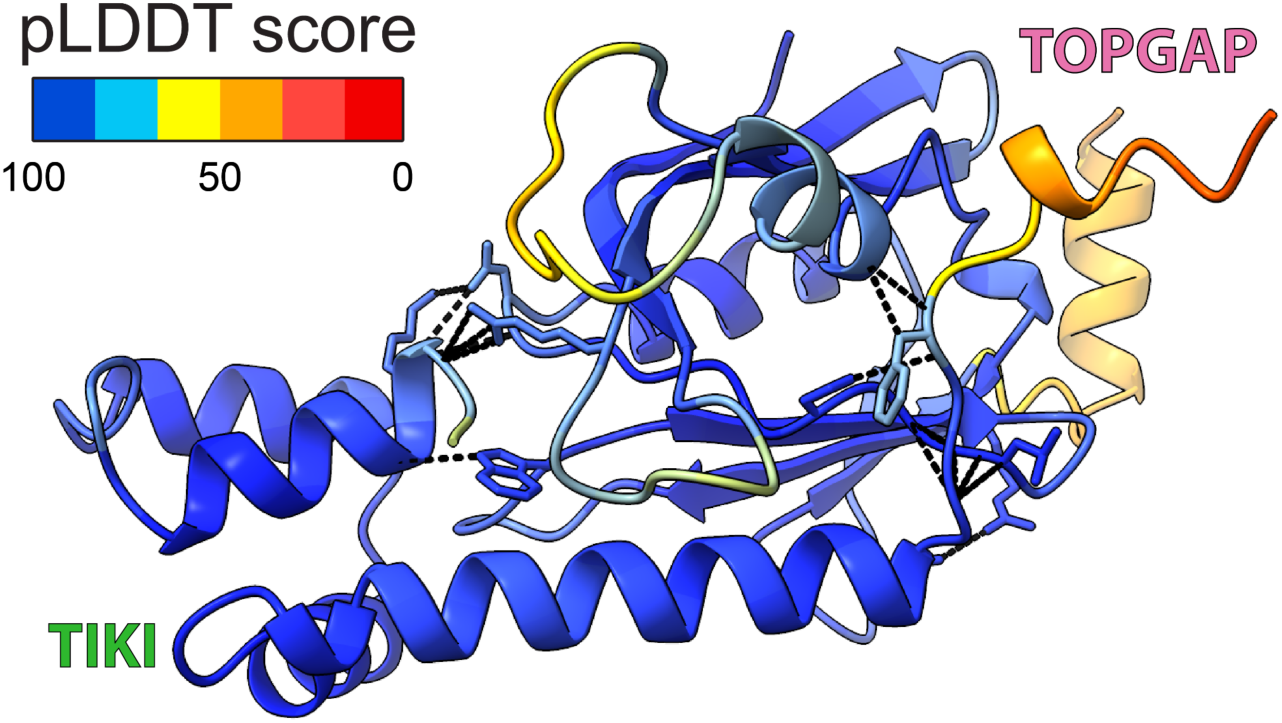
AF2-M–predicted model of TIKI in complex with the RBD of TOPGAP. The TIKI–RBD structural model is colored according to the AlphaFold prediction confidence (pLDDT) scores, as indicated in the rectangular box.

**Figure S11.**
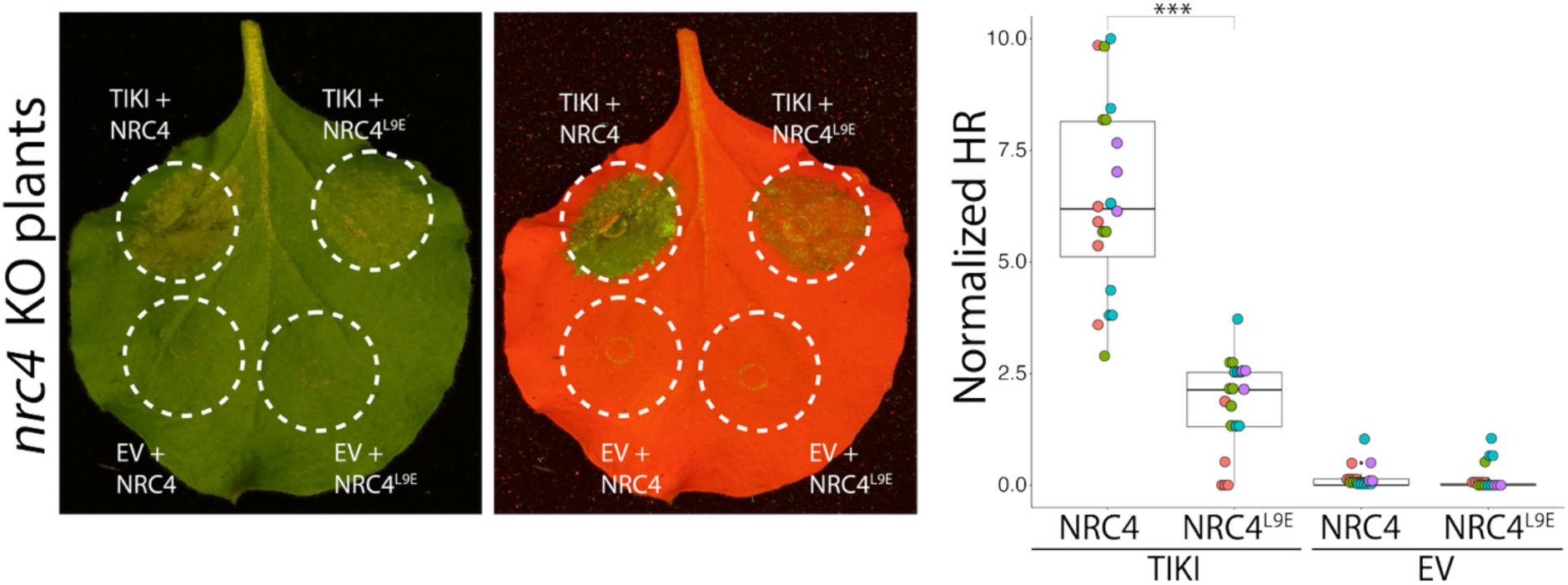
An intact MADA motif in NRC4 supports robust TIKI-triggered hypersensitive response. Representative images of *nrc4* KO *N. benthamiana* leaves infiltrated with TIKI or EV control together with wild-type NRC4 or the MADA-motif mutant NRC4^L9E^. Daylight and UV images were captured at 3 dpi and HR was quantified. Reintroduction of wild-type NRC4 restores robust TIKI-triggered HR in *nrc4* KO plants, whereas expression of the NRC4^L9E^ mutant fails to restore robust HR, resulting in significantly reduced cell death. Box-and-dot plot showing that WT NRC4 significantly enhances TIKI-triggered HR (6.45, n = 20) compared with NRC4^L9E^ (1.83, n = 20). Expression of NRC4 or NRC4^L9E^ together with EV does not induce visible HR. Statistical comparisons were performed using Mann–Whitney U tests with Bonferroni correction in R. ***P < 0.001.

